# Using genomes and evolutionary analyses to screen for host-specificity and positive selection in the plant pathogen *Xylella fastidiosa*

**DOI:** 10.1101/2022.04.25.489460

**Authors:** Tiffany N. Batarseh, Abraham Morales-Cruz, Brian Ingel, M. Caroline Roper, Brandon S. Gaut

**Affiliations:** Department of Ecology and Evolutionary Biology, UC Irvine, Irvine, CA, USA 92697; Department of Plant Pathology, UC Riverside, Riverside, CA 92502

**Keywords:** Evolution, virulence, genomics, phylogenetic analysis, positive selection

## Abstract

*Xylella fastidiosa* infects several economically important crops in the Americas, and it also recently emerged in Europe. Here, using a set of *Xylella* genomes reflective of the genus-wide diversity, we performed a pan-genome analysis based on both core and accessory genes, for two purposes: i) to test associations between genetic divergence and plant host species and ii) to identify positively selected genes that are potentially involved in arms-race dynamics. For the former, tests yielded significant evidence for specialization of *X. fastidiosa* to plant host species. This observation contributes to a growing literature suggesting that the phylogenetic history of *X. fastidiosa* lineages affects host range. For the latter, our analyses uncovered evidence of positive selection across codons for 5.3% (67 of 1,257) of core genes and 5.4% (201 of 3,691) of accessory genes; these genes are candidates to encode interacting factors with plant and insect hosts. Most of these genes had unknown functions, but we identified some tractable candidates including *nagZ_2*, which encodes a beta-glucosidase that is important for *Neisseria gonorrhoeae* biofilm formation; *cya*, which modulates gene expression in pathogenic bacteria; and *barA*, a membrane associated histidine kinase that has roles in cell division, metabolism, and pili formation.

**ABSTRACT IMPORTANCE:** *Xylella fastidiosa* causes devasting diseases to several critical crops. Because *X. fastidiosa* colonizes and infects many plant species, it is important to understand whether the genome of *X. fastidiosa* has genetic determinants that underlie specialization to specific host plants. We analyzed genome sequences of *X. fastidiosa* to investigate evolutionary relationships and to test for evidence of positive selection on specific genes. We found a significant signal between genome diversity and host plants, consistent with bacterial specialization to specific plant hosts. By screening for positive selection, we identified both core and accessory genes that may affect pathogenicity, including genes involved in biofilm formation.

## INTRODUCTION

Bacteria exhibit extensive intraspecific variation in genome content. This variation is the raw material for evolutionary adaptation, including the evolution of pathogenicity and virulence (1–4). One example of genome variation comes from an early study of *Escherichia coli* that compared two pathogenic strains and one non-pathogenic laboratory strain (5). Of the entire set of protein coding genes annotated by the three genomes, only 39.2% were shared among the three isolates. Intriguingly, the two pathogenic strains each had 1,300 unique genes, while the laboratory strain had only 585, suggesting that genes that vary across accessions (i.e., accessory genes) contribute to virulence. Similar patterns have been illustrated for plant pathogens (6, 7). In *Xanthomonas*, for example, horizontal gene transfer (HGT) has shuffled virulent accessory genes from pathogenic strains to previously non-pathogenic strains (4), facilitating the infection of common bean (*Phaseolus vulgaris* L.). In short, accessory genes contribute to host-pathogen interactions, making them a critical focus for comparative analyses of genome evolution and function.

Here we investigate variation in the genome content of another plant pathogen. *Xylella fastidiosa* is endemic to the Americas and was first identified as the causal agent of Pierce’s Disease (PD), an economically devastating disease in grapevines (*Vitis vinifera* ssp. *vinifera*) (8, 9). *X fastidiosa* causes additional economically and ecologically impactful diseases like citrus variegated chlorosis, coffee leaf scorch, oak leaf scorch and elm leaf scorch, among others. Historically, the geographic distribution of *X. fastidiosa* was limited to the Americas, but it was recently introduced to the European continent by anthropogenic transmission, which has further expanded its host range and led to emerging diseases like olive quick decline syndrome (OQDS) in Italy (10, 11). *X. fastidiosa* has since been detected in various plants species across locations in Europe including France, Spain, and Portugal (12, 13). In susceptible hosts, *X. fastidiosa* can lead to significant crop losses, and it continues to threaten crops globally (14, 15).

For each of these diseases, *X. fastidiosa* is transmitted by xylem-feeding insect vectors into the plant host, where it then utilizes cell wall degrading enzymes to systemically colonize the xylem. In the xylem, it forms biofilms that are thought to be integral to pathogenicity (16, 17). Colonization is also governed, in part, by virulence and pathogenicity factors that influence a wide range of bacterial functions – e.g., biofilm formation, host cell wall degradation, regulatory systems, stress responses and bacterial membrane composition -- although other abiotic factors (like plant drought stress) likely also contribute to disease progression (13). Given its economic impact, the effects and mechanisms of *X. fastidiosa* infection have been studied widely, especially in grapevine (18). However, many pathogenicity factors likely remain undiscovered, and crucial questions remain unanswered about the genetic factors that govern host-pathogen interactions and potential host specialization (13).

In this context, it is helpful to recognize that *X. fastidiosa* consists of three commonly recognized subspecies that form distinct phylogenetic clades: ssp. *fastidiosa*, *multiplex*, and *pauca.* Each subspecies has unique phenotypic characteristics and DNA markers (19). Two other subspecies, *morus* and *sandyi*, have also been suggested, though they are not recognized as broadly (9); indeed, *morus* is believed to be a product of a recombination event between *fastidiosa* and *multiplex* isolates (8). The recognition of subspecies is critical, because initial work suggested that subspecies correlate with specific plant hosts (20). While it has long been known that that genetic differences among strains facilitate host-plant specialization (18, 21–23), there is not a clear one-to-one correspondence between pathogen and host. For example, some strains can infect more than one host species, as demonstrated by a strain that causes PD in grapevines and also leaf scorch in almonds (21). Consequently, the questions of the evolution of and determinants of host specificity are still central for understanding the distribution and effects of this pathogen.

In this study, we analyze *X. fastidiosa* genome evolution among isolates from different plant hosts. Our study is not unique in some respects, because numerous comparative genomic studies of *X. fastidiosa* have been published already. Many of these studies have focused on clarifying phylogenetic relationships. For example, Marcelletti and Scortichini (2016) studied 21 genomes to resolve taxonomic relationships among subspecies; Giampetruzzi et al. 2017 extended sampling to 27 genomes, in part to place a novel strain (ST53) in the broader *X. fastidiosa* phylogeny; and Denancé et al. (2019) used kmers from 46 genomes to untangle species and subspecies relationships. Another recent study compared *X. fastidiosa* populations from Central/South America (Costa Rica, Brazil), North America (California, Southeastern US), Europe (Spain, Italy), and Asia (Taiwan) to elucidate the evolutionary origins of the subsp. *fastidiosa* and *pauca* (24). Still other studies have focused on populations. For example, Vanhove et al. (2020) isolated and sequenced *X. fastidiosa* subsp. *fastidiosa* from symptomatic grapevines from five different California locations (25).

One common theme of genomic studies is that they identify the set of genes that are present in most samples (i.e., core genes) and used those genes as the basis to perform phylogenetic inference. These phylogenies have been used for various purposes. For example, two recent papers have used phylogenies to explore the question of host specificity. In one, Uceda-Campos et al. (2022) found that *X. fastidiosa* isolates grouped on the phylogeny by geography, but not by plant host species, suggesting host specificity is not correlated with phylogenetic relationships and genetic divergence (26). In contrast, Kahn and Almeida (2022) used the phylogeny to infer the ancestral character states of plant hosts and found that the ancestral host plant could be inferred for most ancestral nodes (27). They concluded that genetic history affects host range and also identified ∼30 genes whose presence/absence correlated with specific plant hosts.

In this study, we combined 20 new *X. fastidiosa* genomes with publicly available data to build a dataset for molecular evolutionary analysis and to investigate patterns of host specificity in a phylogenetic context. For host-specificity analyses, we focused on core genes, but we also assessed the phylogenetic signal, pattern of gene gain and loss, and potential host associations of accessory (i.e., non-core) genes. Our goals for these analyses were to add to the growing literature about genetic correlations between phylogenetic history and host specificity but also to further consider the dynamic evolution of accessory genes in this context (27). In addition, we performed extensive analyses of the ratio of nonsynonymous to synonymous (dN/dS or ω) substitutions to identify genes under positive selection (ω > 1.0). Genes under positive selection may be involved in arms-race (or Red-Queen) dynamics between pathogens and hosts (28, 29). In other systems, ω analyses have identified genes with functions that contribute to host defense and also discovered entirely new sets of genes and pathways involved in pathogen-host interactions (30–32). Here we apply tests for positive selection in the hope of gaining insights into the sets of genes that may affects host-pathogen interactions.

## METHODS

### Novel X. fastidiosa genomes

Fully extracted DNA from 20 *X. fastidiosa* isolates were provided by the French Collection of Plant-Associated Bacteria (CIRM-CFBP; http://www6.inra.fr/cirm_eng/CFBP-Plant-Associated-Bacteria) and from the University of California, Riverside. Genomic DNA was prepared for Illumina sequencing using the Illumina Nextera DNA Flex Library Prep kit, following the manufacturer’s recommendations and for Pacific Biosciences (PacBio) sequencing with the SMRTbell Express Template Prep Kit 2.0. SMRTbell libraries had 10kb DNA target insert size (Pacific BioSciences, Menlo Park, CA) using 360ng of sheared DNA as input. DNA libraries were sequenced with both Illumina and PacBio technologies at the University of California, Irvine Genomics High Throughput Facility (https://ghtf.biochem.uci.edu). The Illumina sequencing reads were quality assessed using FastQC, and reads were trimmed using Trimmomatic v. 0.32 (33, 34) using default options. PacBio sequencing reads were corrected and trimmed using Canu v. 1.5 (35). The long and short reads were used for genome assembly with Unicycler v. 0.4.8 in hybrid assembly mode (36). Genome assembly statistics were calculated using Quast v. 5.0.2 (37). As is common practice (38), short contigs (<500 bp) were removed from the assemblies using Seqkit v. 0.13.2 (39).

### Genome Assembly of public data and sample set curation

We complemented our set of novel genomes with publicly available data. To do so, we downloaded all available whole genome assemblies of *X. fastidiosa* and *X. taiwanensis* (as an outgroup) from the National Center for Biotechnology Information (NCBI) and Sequence Read Archive (SRA) databases on July 9, 2020 (Supplementary Table 1). In addition, we downloaded the raw, short-read sequences for an additional 20 isolates (25, 40). For each isolate, we gathered information about its geographic origin and its host plant from NCBI and from the Pathosystems Resource Integration Center (PATRIC) database. To assemble the raw reads from the 20 unassembled accessions into genomes, we assessed quality and trimmed the reads and applied SPAdes v. 3.14.0 (41) with the *--careful* option, following Vanhove et al. (Supplementary Table 2; 2020). If long reads were also available, as they were for 5 isolates from the work of Castillo et al. 2020, then whole genome assembly was performed with Unicycler v. 0.4.8 in hybrid assembly mode (36).

In total, we gathered and generated 148 *Xylella* genome assemblies. From this set, we removed isolates that did not have information about their host isolation source or were lab-derived recombinant strains. The remaining 129 genomes were re-annotated by the same method - based on Prokka v. 1.14.6 analysis – that we applied to the new genomes, to ensure homogeneity. The Prokka analyses were then input into Roary v. 3.13.0 with options *-i 80 -cd 100 -e -n -z* to obtain a core gene alignment for initial comparisons among isolates (42, 43); we defined core genes as those that were detectable in 100% of the samples. This core set was aligned with MAFFT and polished using gBlocks v. 0.91b (44–46). The polished alignment was used as input for RAxML v. 8.2.12 to build a preliminary phylogenetic tree (47), which we used to evaluate and curate the isolates (Supplementary Figure 1).

To curate the dataset, we created a distance matrix from the RAxML phylogenetic tree, using the Tree and reticulogram REConstruction (T-REX) server (48). Many of the genomes – most of which were gathered for population genomic analyses - were sampled from the same plant host and were nearly identical genetically. To limit sampling biases for our species-wide study, we removed clones and near-clones based on the distance matrix. That is, if two or more isolates had a pairwise distance ≤ 0.0001 and came from the same host, we retained the isolate with the more contiguous assembly. We also used CheckM (49) to assess genome completeness based on a set of conserved single copy genes (Supplementary Table 3). After applying these filters, our final dataset consisted of 63 *X. fastidiosa* genomes and one outgroup genome (*X. taiwanensis* PLS229) that were isolated from 23 distinct plant host species (Supplementary Table 1).

### Pan-genome analysis

To perform a pan-genome analysis, we applied Roary to the 64 *Xylella* genomes using *gff* files from Prokka as input. Roary was applied with the option *-i 80,* as used in previous microbial studies (38, 50), to lower the blastp sequence identity to 80% from the default 95%. We defined a core gene as a gene present in 95% of the isolates used in the analysis (i.e., a core gene was present in at least 60 of 63 *X. fastidiosa* accessions). From the Roary output, we extracted a representative nucleotide sequence of each core and accessory gene using cdbfasta (https://github.com/gpertea/cdbfasta) and translated the nucleotide sequence to amino acids using the transeq command from EMBL-EBI (51). The representative sequences were the basis for functional categorization -- using the eggNOG-mapper v. 2 (52, 53) -- of both core and accessory genes. Gene Ontology (GO) enrichment analyses were performed online at (http://geneontology.org) using *Xanthomonas campestris* as the reference list (54). To explore function further, we also used the Conserved Domain Database online tool (https://www.ncbi.nlm.nih.gov/cdd/) to identify protein domains.

### Phylogenetic Tree Construction

We used the core gene alignment from Roary to build a phylogenetic tree, based on a subset of genes that were present in all 63 *X. fastidiosa* samples and the *X. taiwenensis* outgroup. To do so, we curated the alignments with gBlocks v. 0.91b (44), used the polished alignment as input for IQtree v. 2.0.3, and selected the best nucleotide model for phylogenetic tree construction (55, 56). We ultimately constructed an unrooted tree using the GTR+F+R8 model with RAxML (Stamatakis 2014), using the ‘best tree’ option. Phylogenetic trees were visualized and annotated using the ape package v. 5.5 in R v. 4.0.2 (57, 58). We used the most likely phylogeny to test associations between phylogenetic relatedness, geography, and host isolation source (plant taxonomic order information taken from https://www.itis.gov/) with ANOSIM implemented in the vegan package v. 2.5-7 in R (59).

*X. fastidiosa* is naturally transformant and undergoes homologous recombination (9, 60), but recombined genomic regions can obscure vertical phylogenetic relationships. To account for potential recombination among *X. fastidiosa* genomes, we applied Gubbins v. 3.2.1 (61, 62), using again the subset of genes that were found in all 64 samples. From this input, Gubbins identified regions that were likely to have undergone recombination and removed them from the alignment. We then built a phylogeny from this recombination-adjusted core gene alignment using RAxML, as described above. We assessed the congruence between the two phylogenetic trees (i.e., with and without removal of potentially recombining regions) using phytools v. 1.0-1 in R (63).

Finally, we also built a Neighbor-Joining (NJ) tree based on the presence-absence matrix of accessory genes. We first calculated the Euclidean distances from the presence-absence matrix of the accessory genes using the *dist* function in R (64). We then built an NJ tree from the Euclidean distances using the ape package in R (57). We also utilized the ANOSIM and Mantel test (in the *vegan* package) to measure the correlation between accessory gene content and phylogenetic relatedness. The Mantel test required two distance matrices, which were the Euclidean distances estimated from the accessory gene presence-absence matrix and the distances from the RAxML core gene phylogeny generated by the reticulogram REConstruction (T-REX) server (48).

### Gain and Loss of Accessory Genes

We utilized GLOOME to investigate gene gain and loss dynamics along the core phylogenetic tree of *X. fastidiosa* (68). GLOOME uses a mixture-model approach, coupled with maximum-likelihood inference, to infer rates of gain and loss of genes along the branches of a phylogeny. It takes as input the phylogenetic topology, in this case the phylogenetic topology based on core genes, and a presence-absence matrix of genes. The pattern of genic presence and absence was obtained through M1CR0B1AL1Z3R, as recommended by the GLOOME authors, and then directly input into GLOOME using default settings (69). The default settings included a fixed rate of gene gains and losses with gamma distributed rates across genes (or sites). Among the output, GLOOME returned two phylogenetic trees with branch lengths representing either the number of expected gain events or the number loss events on each branch. As recommended (68), branch lengths representing relative gain and loss rates were extracted from the phylogenetic trees using FigTree v. 1.4.4 (http://tree.bio.ed.ac.uk/software/figtree/). To normalize expected gain (or loss) events with sequence divergence, we calculated the ratio of inferred gain (or loss) against the branch lengths of the sequence-based core phylogeny. Outlier branches with excess normalized gains or losses were identified using the interquartile range criterion.

### Positive selection analyses

We employed codeml from PAML v. 4.9 to calculate ω, the ratio of nonsynonymous to synonymous rates (70, 71). We performed *codeml* analysis on nucleotide alignments of the single-copy core genes, single-copy accessory genes, and multicopy genes (defined as genes with two or more copies in a single accession). For all tests, we required at least four sequences, as the minimum number suggested for *codeml* analysis (http://abacus.gene.ucl.ac.uk/software/pamlFAQs.pdf). For each gene and sequence set, we ran analyses by generating an unrooted maximum-likelihood tree for each gene based on the DNA alignment, using RAxML v. 8.2.12. This approach recognizes that the phylogeny of a single gene may not follow the consensus phylogeny due to a history of recombination. For completeness, however, we also performed codeml analyses by assuming the global phylogeny for the subset of genes that were present in all 64 samples. The outcomes of the two approaches were highly correlated (Supplementary Figure 2), and so for simplicity we focused on results based on phylogenies inferred separately for each gene.

Given the input phylogenies, we performed *codeml* analyses that relied on calculating likelihood ratios (LRs) under various models (Yang 2007). Briefly, we used the models to test the null hypothesis that ω = 1.0 against the alternative of positive selection (ω > 1.0) in two different ways. The first was a global test across the entirely phylogeny of a gene – i.e., across all branches and all sites. This test requires the comparison of two models: one (Model = 0 with Fix_omega = 1 and Omega = 1 in the *codeml* control file) that estimates a single ω from the data and another that sets ω=1.0 (Model = 0 with Fix_omega = 0 in the *codeml* control file). The two models yielded evidence for positive selection when the initial ω estimate was >1.0 and when the likelihood of the two models differed significantly, based on P < 0.01 after FDR correction. The second set of analyses was across sites – i.e., testing for genes with variable selection pressure across sites. For each gene, we first compared models M0 and M3 to test for heterogeneity in evolutionary rates across codons. If that test was significant, we then compared sites models M1a and M2a from *codeml* to test for specific codons with evidence of positive selection (ω > 1.0). For all summary statistics of ω, we excluded estimates of ω that were greater than 10 as potentially unreliable due either to low d_s_ or poorly resolved optimization. Individual codon residues under positive selection were identified using the Empirical Bayes analysis in codeml.

## DATA AVAILABILITY STATEMENT

All high-throughput sequence data generated in this study have been submitted to the NCBI Sequence Read Archive database at https://www.ncbi.nlm.nih.gov/sra and can be accessed with project number PRJNA833428.

## RESULTS

### Core and Accessory Genes in *Xylella*

To investigate genome evolution in *X. fastidiosa*, we sequenced 20 novel *X. fastidiosa* genomes using hybrid approaches and retrieved publicly available genomes and raw sequencing data (Supplementary Tables 1 and 2). After filtering for isolation source and genetic distance, we retained a sample of 63 genomes that were broadly distributed among the subspecies. All our analyses were performed on this final set of 63 *X. fastidiosa* genomes with the *X. taiwanensis* outgroup. The *X. fastidiosa* genomes ranged in size from 2.42 Mb to 2.96 Mb, with an average length of 2.61 Mb (Figure 1A) and an average of 2,478 predicted genes (Figure 1B). The samples were extracted from 22 plant hosts representing 12 botanical orders (Figure 1C).

**Figure 1:**
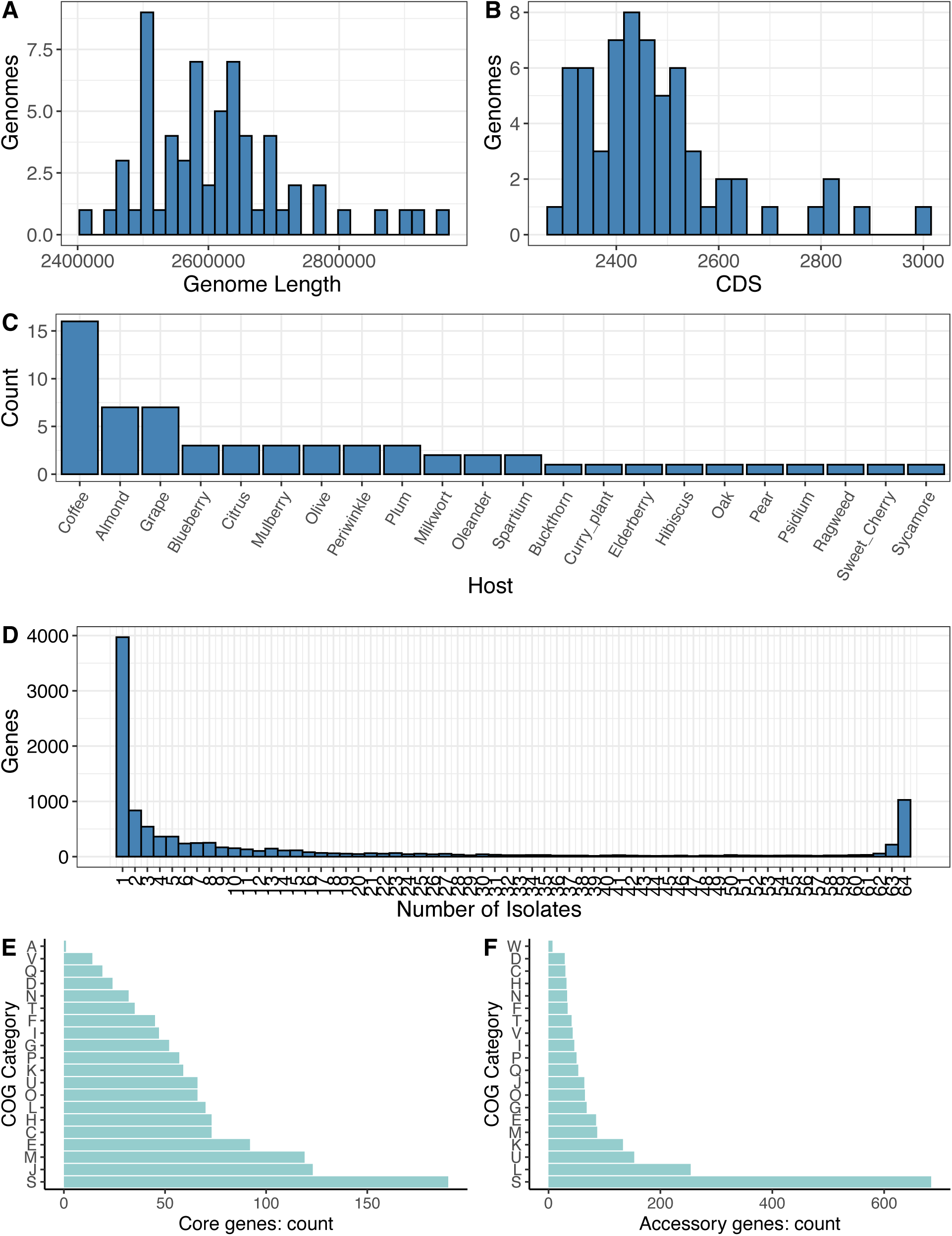
Histograms reporting the characteristics of the 64 *Xylella* genomes. A) Genome lengths, exhibited in base pairs. B) The number of genes within a genome. C) A histogram of the plant species from which genomes were isolated. D) A histogram of the number of genes found in *x* number of genomes; this histogram shows, for example, that nearly 4,000 genes are found in only of one the genomes out of the entire sample of 64 genomes and that 1,024 genes are found in all 64 genomes; E) the distribution of functional categories for the set of 1,257 core genes and F) the distribution of functional categories for the set of 9,220 accessory genes. A key to the COG categories for panels E) and F) is in Figure S4.

We categorized each gene as either core (present in 95% or more of *X. fastidiosa* samples) or accessory (43). Across all 64 genomes, we identified 10,477 genes within the pan-genome; of those, 1,257 were core genes and 9,220 were accessory genes, with nearly 4,000 genes found in only a single isolate (Supplementary Table 4; Figure 1D). We performed functional analyses on both the core and accessory gene sets by grouping protein coding sequences into clusters of orthologous gene (COG) (Figure 1E-F). We compared COG category rankings between the core and accessory gene sets, with significant differences (Wilcoxon rank sum test, paired; P = 0.0001). After excluding genes with unknown function, the largest COG categories in the core gene set were ‘translation, ribosomal structure, and biogenesis’ (123 genes), ‘cell wall/membrane/envelope biogenesis’ (119 genes), and ‘amino acid transport and metabolism’ (92 genes). In contrast, the largest categories for accessory genes were ‘replication, recombination and repair’ (547 genes), ‘intracellular trafficking, secretion, and vesicular transport’ (364 genes), and ‘transcription’ (298 genes). Additionally, we investigated the core and accessory gene lists for significant GO based enrichment of specific biological processes (Supplementary Tables 5 and 6).

### Phylogenetic Patterns of Core Genes, Accessory Genes and Hosts

#### Phylogenetic Relationships

We constructed a maximum likelihood phylogeny based on a subset of 1,024 genes that were present in all 64 isolates. The topology was highly supported; it had a mean bootstrap support of 93.75% across all nodes, with a median of 100% (Figure 2). The lowest bootstrap supports were primarily found at nodes separating *X. fastidiosa* strains isolated predominantly from grapevines, reflecting relatively low evolutionary divergence among these samples. As expected (72), isolates clustered into three distinct clades representing the three main subspecies (ssps. *fastidiosa*, *multiplex*, and *pauca*), with 27, 23 and 13 isolates in each clade, respectively. To account for the possibility that homologous recombination impacted the resolution of the core phylogeny, we extracted regions of the core gene alignment that had an apparent history of recombination (61), ultimately removing 85.1% of the alignment. The phylogeny inferred from this alignment was nonetheless highly congruent with the phylogeny that did not consider recombination. Only five accessions had altered positions between the recombination-adjusted and non-adjusted trees (Supplementary Figure 3).

**Figure 2:**
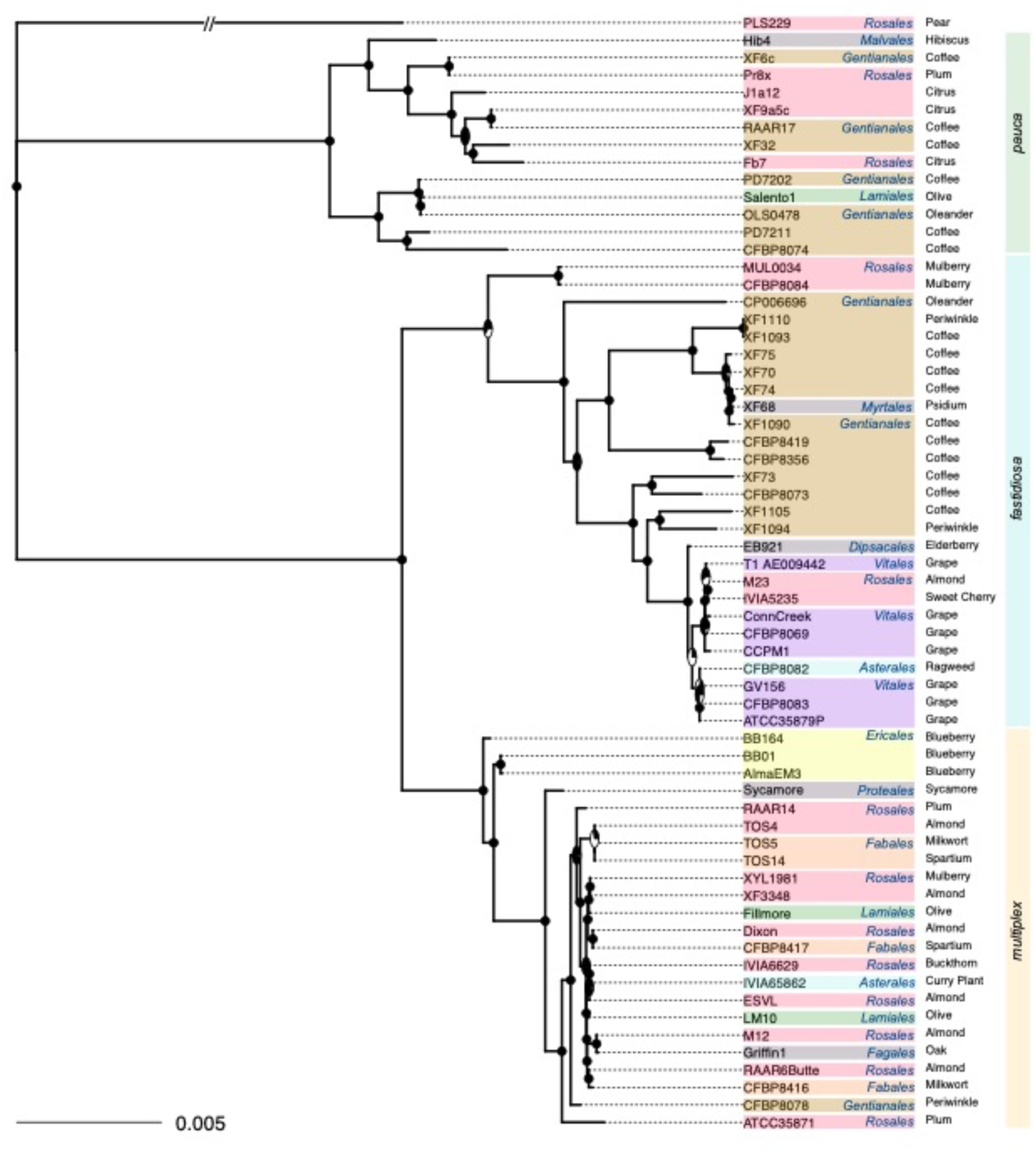
The inferred phylogeny of 64 *Xylella* genomes, based on maximum likelihood inference of core gene alignments. Each isolate is labelled at the tips and is colored according to the order of the plant isolation source (host). The common name of the host is provided to the right of order information. The three *X. fastidiosa* subspecies are indicated, as are bootstrap values at each node. The bootstrap values are pie charts, where black represents the percent of bootstrap support, and the scale bar reflects the magnitude of sequence divergence per nucleotide site.

To investigate general evolutionary patterns of the accessory gene complement, we compared the core gene phylogeny against a phylogeny based on accessory gene composition (Figure 3). Both the core gene and the accessory gene phylogenies clustered into three groups, and all members of the groups were consistent between phylogenetic treatments. This pattern broadly suggests that accessory genes, while defined by their inconstancy, are not exchanged *en masse* to a sufficient degree to alter phylogenetic signal among subspecies. Within subspecies, however, relationships at the tips of the phylogeny often differed between core and accessory trees. As an example, the cluster corresponding to *multiplex* displayed the most discordance between the core and accessory trees, with all OTUs contributing to phylogenetic incongruence (Figure 3). Interestingly, our *multiplex* sample also had more plant host species than our *fastidiosa* and *pauca* samples, suggesting the possibility (but by no means proving) that host factors may affect or moderate genome content (27). Nonetheless, we found a significant correlation between distance matrices based on the core and accessory phylogenies (Mantel test, R = 0.1144, P = 0.019), which is consistent with the fact that the two trees had the same three major clades. The overarching impression of these analyses is that accessory gene composition does not turn-over so rapidly, due to HGT or other mechanisms, to erase phylogenetic and historical signals of subspecies diversification within *X. fastidiosa*.

**Figure 3:**
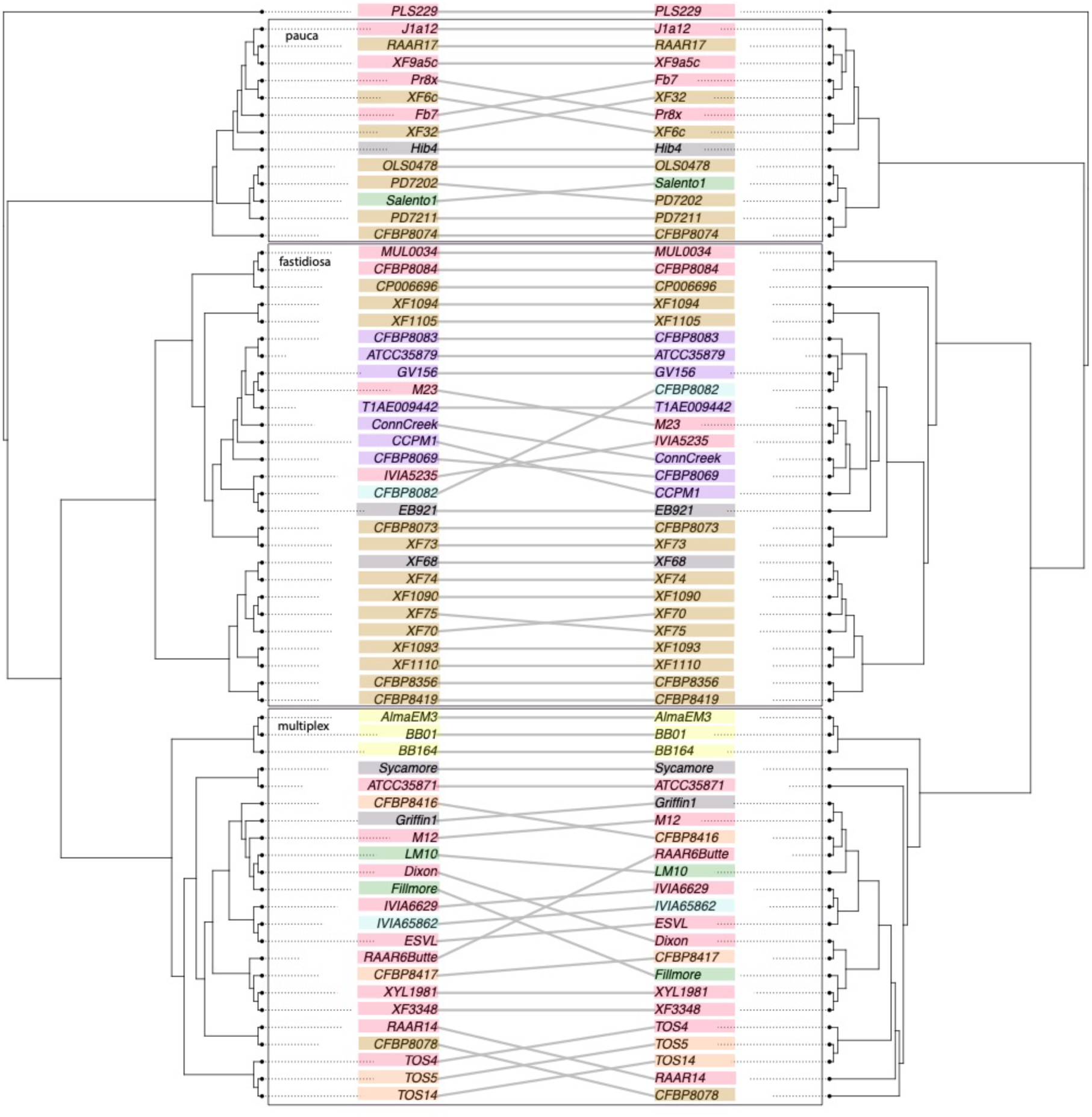
A comparison of a NJ tree based on distances due to gene presence / absence (on the left) to the likelihood tree based on the core gene alignments (from Figure 2, on the right). As in Figure 2, the isolates are labelled at the tips of trees, with the colors representing plant order. Both phylogenies contain three main *X. fastidiosa* clades, representing the three subspecies. Lines connect the same isolate between the two trees, with angled lines representing topological discordance between phylogenies. The three *Xylella* subspecies are outlined in a black box and labelled.

We used both species phylogenies (based on alignments with and without putative recombinant regions) to test for associations between *X. fastidiosa* and their isolation sources (i.e., geographic location or host plant information) using ANOSIM (see Methods). There was a weakly significant phylogenetic association (ANOSIM R = 0.08178, P = 0.042) between the geographic location and the phylogeny built from the full core gene alignment (ANOSIM R = 0.08178, P = 0.042) but not with the phylogeny built from non-recombinant regions (ANOSIM R = −0.004147, P = 0.4895). Applying the same approach to host species revealed a significant phylogenetic signal for both phylogenies (ANOSIM R = 0.1381, P = 0.047; non-recombining regions only, ANOSIM R = 0.6698, P < 1 x 10^-4^). Since *X. fastidiosa* infects a wide range of plants, we also retrieved the taxonomic order of each plant host to test for a phylogenetic signal at a deeper taxonomic level, recapitulating the significant association with both phylogenies (ANOSIM R = 0.3152, P < 0.0001; non-recombining regions only, ANOSIM R = 0.1226, P = 0.0198). In other words, strains isolated from plants within the same taxonomic order were more phylogenetically similar to one another than isolates taken from unrelated plants.

We hypothesized that accessory genes are crucial in pathogen-host interaction and therefore repeated ANOSIM analyses with a distance matrix based on the presence and absence of accessory genes (Figure 3). We found a significant association between accessory gene content and geographic isolation source (ANOSIM R = 0.4553, P = 0.5307) and a weakly significant association between accessory gene content and host species (ANOSIM R = 0.1503, P = 0.0372). The association was lost, however, at the level of plant order (ANOSIM R = 0.02367, P = 0.3033). Overall, associations were less evident based on accessory gene content vs. the core-gene phylogeny.

#### Gene Gain and Loss

The sheer number of accessory genes indicate that the genome content of *X. fastidiosa* is, like other microbes (73, 74), shaped by extensive gene gain and loss events that are probably mediated by HGT (75). We were interested in assessing the pattern of gene gain and loss across the phylogenetic tree, hypothesizing that both could be enhanced on branches that lead to host shifts. We used GLOOME to estimate the number of gain and losses of accessory genes across the *X. fastidiosa* phylogeny and represented those estimates phylogenetically (Figure 4). Ignoring the branch leading to the *X. taiwanensis* outgroup (PLS229), the internal branches discriminating the *X. fastidiosa* subspecies were estimated to average ∼550 separate gene gain and gene loss events. The remainder of the tip and ingroup branches averaged ∼100 gene gain and loss events (average gains/branch = 92.8 genes; average losses/branch = 100.0 genes; Figure 4A&B).

**Figure 4:**
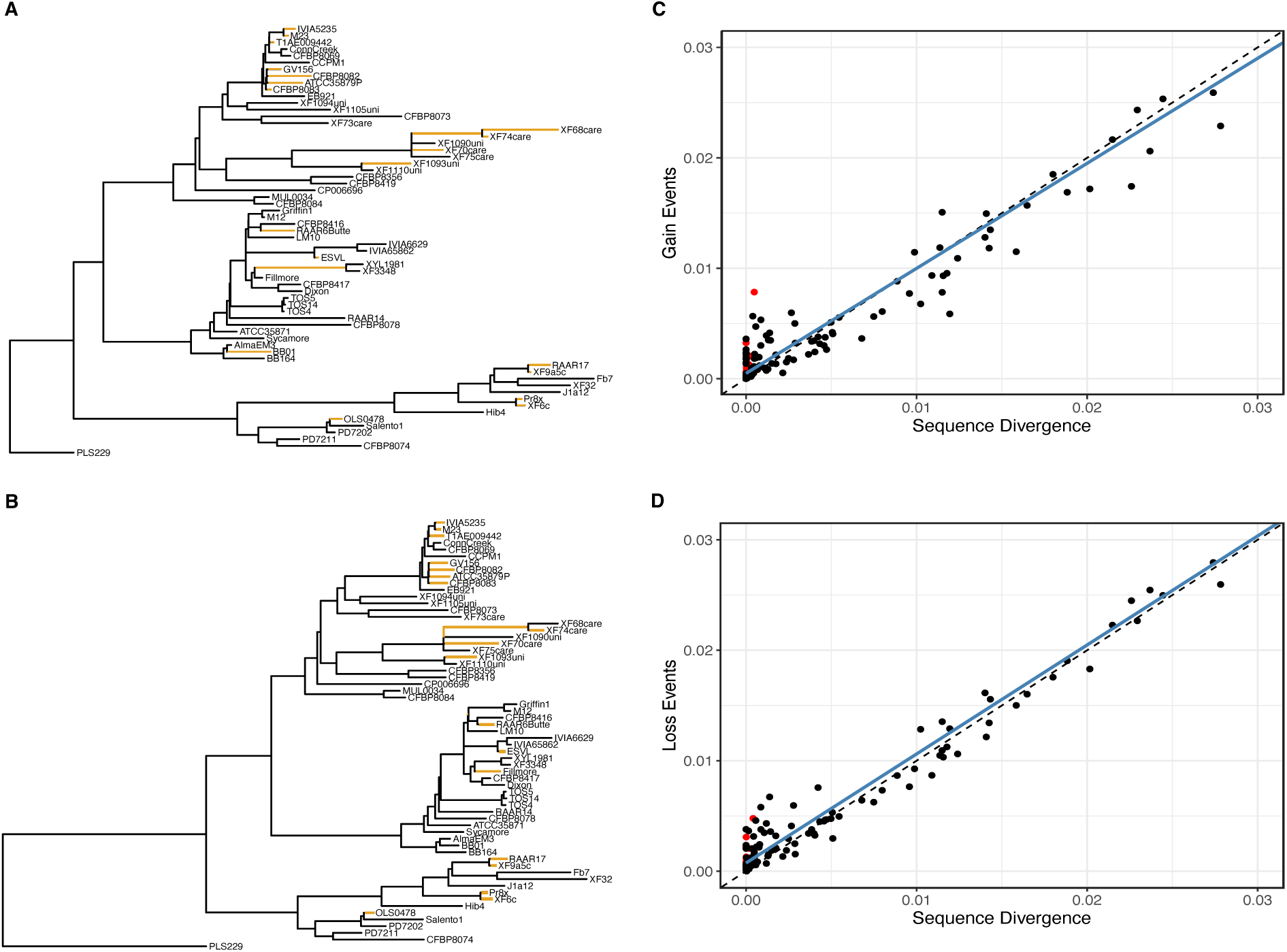
The result of gene gain and loss analyses. A) The phylogeny of the isolates, with branch lengths proportional to the number of gene gain events. The colored branches are branches with outlier gene gain rates. B) The phylogeny of the isolates, with branch lengths proportional to the number of gene loss events. The colored branches are branches with outlier gene loss rates. C) A plot of the gene gains against sequence divergence; in the plot each dot represents one of the 125 branches on the phylogeny. Outlier dots are colored red. D) As in C, with gene losses plotted again sequence divergence.

While it is useful to estimate the number of gains and losses on each branch, we thought it more helpful to normalize the number of estimated gain and loss events by branch lengths, estimated from the sequence analysis of core genes. This normalization by branch length converted the number of gene gains and losses to *rates* of gene gain (or loss) relative to sequence divergence. We then sought to identify branches with aberrantly high rates of gene gain or loss (Figure 4C&D), which reflect branches with especially notable turnover of accessory genes. Like a previous microbial study (76), we found that most of the phylogenetic lineages with outlier rates were located at the tips of the phylogenetic tree. For example, of the 21 branches with high rates of gene gain, 19 were at the tips of the phylogeny (Figure 4A). Similarly, 18 of 21 branches with high rates of gene loss were external branches. These observations suggest features about the evolutionary dynamics of genic turnover (see Discussion).

### Characterizing selection with *ω*

We characterized selection on individual genes by estimating the dN/dS ratio (*ω*); we especially sought to identify genes that experienced positive selection (i.e., *ω* > 1.0), as a potential signal of genes that contribute to dynamics between the pathogen and its hosts. To do so, we applied a series of nucleotide substitution models to individual genes, ultimately resulting in tests for positive selection on two levels: globally across a phylogeny and across codon sites (see Methods). For these tests, we examined the full complement of 1,257 core genes, a subset of 3,691 accessory genes, and a set of 187 multicopy genes.

#### Testing selection globally for each gene

We first estimated a single ω value for each gene, using a method that assumes ω is constant across all branches of the entire gene tree and across all codons in the nucleotide alignment. Applied to the core genes, ω estimates (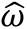) ranged from 0.01048 to 2.92803 with an average of 0.21973 (Figure 5A). Nineteen core genes had 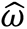 higher than 1.0, but none of these were significantly > 1.0 (P > 0.01, FDR correction). In fact, the vast majority (1,144 of 1,257) of core genes had 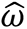 significantly < 1.0 (P < 0.01, FDR correction; Figure 5A), reflecting pervasive purifying selection. The range of 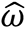 was substantially broader for accessory genes, from 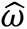 = 0.0001 to 9.60069, with an average of 0.51443 (Figure 5B). Among accessory genes, 367 (9.9%) had a global estimate of ω > 1.0, but only eight had statistically significant evidence for positive selection. These eight genes were candidates to encode proteins involved in host-pathogen interactions, but seven of eight were annotated as hypothetical genes (Supplementary Table 7). Overall, the average 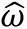 was significantly higher in the accessory gene set compared to the core genes (Welch’s T-test, P < 2.2 x 10^-16^), reflecting either lower purifying selection against these genes, more positive selection, or both.

**Figure 5:**
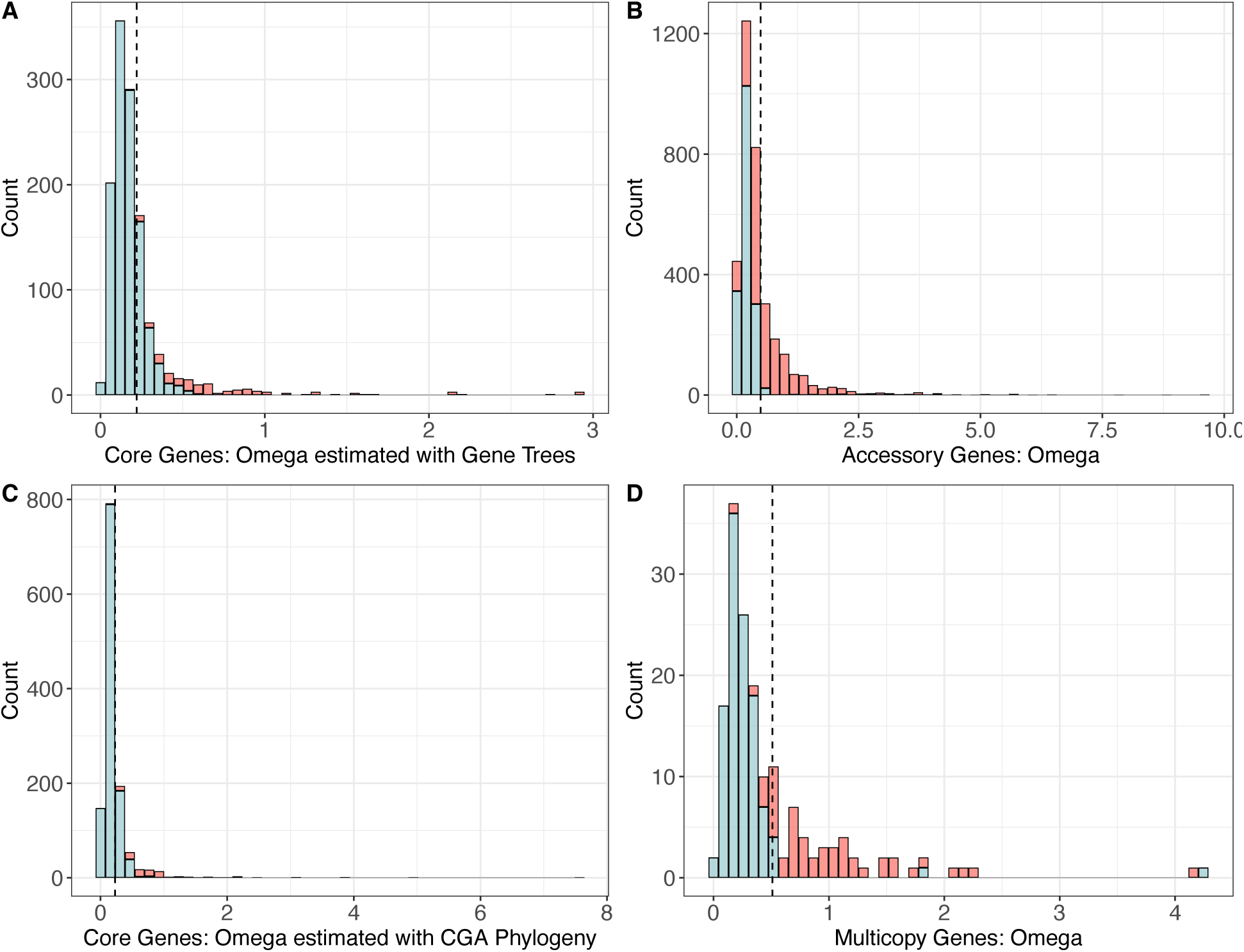
Estimated values of ω under M0 (the one-ratio model) in the core and accessory genes. The distribution of 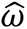 values is plotted for A) core genes estimated with gene trees, B) accessory genes with gene trees C) core genes estimated with the core gene alignment (CGA) phylogeny, and D) multicopy genes with gene trees. Histogram bars are shaded to reflect the outcome of the likelihood ratio test (non-significant tests are colored red and significant tests are colored blue) between a model that estimated 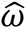 and a model with ω fixed to 1.0. The horizontal dashed line denotes 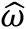 for each gene set.

We also identified 187 genes that had 2 or more copies within a single accession in a syntenic context, but that were single copy in other accessions. We performed *codeml* analysis to estimate ω for each multicopy gene;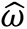 ranged from 0.02272 to 4.26800, with an average of 0.51129 (Figure 5D). Over half of the genes had 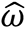 significantly < 1.0 (59.4%; P < 0.01, FDR correction); only one, a hypothetical gene (*group_1109*) had 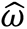 significantly higher than 1.0 (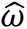 = 1.85845, P < 0.01, FDR correction).

#### Positive selection in codon sites

The global test is a conservative criterion to search of positive selection, perhaps overly so. Accordingly, we turned to an alternative method that tests for variation in ω among codon sites and identifies whether sites are under positive selection. To do so, we ran the sites models in *codeml*, which are a group of nested models. For completeness, we first compared sites model M0, which represents the null hypothesis that there is a single ω value for all sites, against sites model M3, which permits ω to vary among sites. In the core genes, the likelihood ratio test was significant for 501 genes (P < 0.01, FDR correction). We then took this set to compare to test for positive selection using sites models. A total of 67 core genes had evidence of positive selection among sites (P < 0.01, FDR correction). We also tested for positive selection on codon sets within the 3,691 accessory genes using the same approach. Of the total, 895 displayed evidence of variable ω among sites (P < 0.01, FDR correction) and 201 yielded evidence of positive selection (P < 0.01, FDR correction). Finally, we applied the sites models to the set of 187 multicopy genes, yielding another 33 genes with evidence for positive selection. To sum, 5.3% (i.e., 67 of 1,257) of core genes, 5.4% (201/3,691) of accessory genes and 17.6% of multicopy genes had significant evidence of at least one codon with an apparent history of positive selection. Among the 201 accessory genes, four (*cya, group_454, group_1057*, and *group_3542*) also had evidence for positive selection via the global test.

## DISCUSSION

Host-pathogen interactions can drive rapid evolution of pathogenic bacteria, particularly for genes involved in arms-race dynamics (28, 77). Here, we investigated the genomic evolution of the plant pathogen, *X. fastidiosa*, through comparative genomic analysis of genomes that represent diversity across the species, based on a sample set of 64 genomes. The sample was isolated from 23 different plant hosts (Figure 1C) from throughout the world (Supplementary Figure 1). With these data, we constructed a pangenome that contained 1,257 core genes and 9,220 accessory genes, similar to previous studies (40, 78). Of the core genes, the majority were, as expected (79), involved in essential cellular processes -- such as translation, cell wall biogenesis, and amino acid metabolism (Figure 1E). We used the set of core genes to infer a maximum likelihood phylogeny, either with or without adjusting to putatively recombining regions of the genome (Figure 2; Supplementary Figure 3). As with previous systematic treatments of *X. fastidiosa* (19, 72, 80), both phylogenies identified three clades corresponding to the three main subspecies (*fastidiosa, multiplex*, and *pauca*).

We employed both phylogenies to investigate the relationship between the *X. fastidiosa* phylogeny and the plant host. The question of host specialization was first addressed using phylogenetic approaches with multilocus sequencing typing (MLST) data. In this work, Sicard et al. (2018) generated MLST data from 7 housekeeping genes from 50 *X. fastidiosa* genotypes. After building a phylogeny, they tested coevolutionary relationships between host species and *X. fastidiosa* MLST types but found no significant evidence of coevolution, implying a lack of host specialization. This topic was recently revisited with full genome data (26, 27), but the results were inconsistent between studies. Uceda-Campos et al. (2022) found no evidence that plant host species clustered on their *X. fastidiosa* phylogeny, but the samples did cluster by geography. In contrast, Kahn and Almeida (2022) inferred ancestral character states of plant hosts on the *X. fastidiosa* phylogeny and were able to resolve the character state of some deep nodes. They inferred, for example, that coffee plants were the ancestral host species for the node separating *X. fastidiosa* ssp. *fastidiosa* from other subspecies. These patterns suggest that phylogenetic history is associated with specific plant hosts and host ranges.

The disagreement among previous studies, and the fact that all such analyses are properties of the sampled isolates, makes the issue worthy of further assessment. In our study, we have found a significant, non-random association between phylogenetic relationships and both the species and taxonomic order of plant hosts (P < 0.0001) based on core phylogenies. These results are consistent with some level of specialization of *X. fastidiosa* to plant hosts and with the recent analysis of Kahn and Almeida (2022). Moreover, these results were robust to phylogenetic treatment – i.e., the inclusion or exclusion of genomic regions inferred to have histories of recombination. Although it is difficult to quantitatively compare ANOSIM results across studies, it is worth noting that the association of *X. fastidiosa* to plant order is similar in magnitude to the association between a gut colonizing bacterium (*Bifidobacterium*) and the host species from which it was isolated (50).

Given some evidence for host specialization, we hypothesized that it is driven in part by accessory gene content. Under this hypothesis, we predicted an association between genes and hosts should be as (or more) pronounced for accessory genes than for core genes. Instead, we found no significant association between accessory gene complement and taxonomic order and only a weak association with plant species. Our results are unlike, for example, the case of bifidobacteria, where the association with host species was nearly as strong for accessory genes as for host genes (50). We cannot be sure why we do not detect a signal for host specialization of accessory genes, but we can think of three explanations. One is that that host associations, to the extent they exist, are not driven by accessory genes but rather by evolutionary divergence in core genes. Another is statistical power: because there are many more sequence changes among the core genes than there are changes in accessory gene content, the distance matrix for core genes likely has a higher signal-to-noise ratio than accessory gene content. Finally, if accessory genes do mediate host shifts, it is possible – and even likely - that only a subset of accessory genes drive these shifts. Under this scenario, there may be significant associations for a small subset of accessory genes, but the signal of this association is weak across the entire accessory gene set. This conjecture seems reasonable given that Kahn and Almeida (2002) found that the presence/absence of a subset of only ∼30 accessory genes correlated with the plant host. In addition, it is worth emphasizing that *X. fastidiosa* interacts not only with plants but also insect vectors and microbial communities, so that some subset of accessory genes likely contribute to these interactions instead of those with plant hosts.

### The pattern of gene gain and loss events

Another potential tool to study adaptation to specific hosts is by examining shifts in gene composition through gene duplication, deletion, or HGT events (81, 82). We estimated the number of gene loss and gain events along the core-gene phylogeny, and then normalized those numbers relative to sequence divergence. Using this approach, we found that most branches followed a consistent rate of gene gain or loss relative to sequence divergence. The fact that the accessory gene phylogeny recapitulates the three subspecies (Figure 3) suggests, along with previous evidence, that *X. fastidiosa* evolves predominantly through vertical inheritance and intraspecific recombination, rather than HGT from other bacterial species (20, 83).

We have, however, identified 19 and 18 lineages with enriched gain or loss events, respectively, and most of these branches were at the tip of the phylogeny. Again, a potential explanation for these gain and loss dynamics is that they reflect host shifts. There are some isolated examples that are consistent with this hypothesis. For example, isolates XF6c, Pr8x, RAAR17, and OLS0478 in *pauca* have branches with enriched gene gains (Figure 4A). Two of these (OLS0478 and Pr8x) were isolated from oleander and plum, respectively, and are the only isolates associated with those plant hosts in their clades, suggesting a host shift. More globally, however, the evidence for this hypothesis is unconvincing. When we, for example, contrast gene gains between pairs of sister taxa with the same plant host, three of the 16 sister pairs had enriched rates of gene gain. This proportion of enriched branches was not significantly lower than the reminder of the tree (P > 0.05; Fisher’s Exact Test), despite the fact that the sister taxa did not experience a host shift. All of these inferences are of course dependent on our sample and ignore the vector component of the *X. fastidiosa* lifecycle, so there are limitations to our conclusions. At present, however, the evidence for an association between host shifts and enhanced gene gain and loss events is weak.

This leaves unexplained the pattern of enriched rates of gene gain and loss at the tips of the tree. We suspect this pattern is analogous to patterns of mutations in populations, as suggested previously (76). New mutations begin as rare, low frequency variants in single individuals. Eventually most of these mutations are removed by the processes of genetic drift and natural selection, so that there are more new mutations in populations than old mutations. In a phylogenetic context, these new mutations would be evident at the tip of the trees, so it may be reasonable to expect higher effective rates of gene gain and loss in the ‘newest’ phylogenetic branches. This explanation only has credence, however, if the observed gain and loss events are both recent – i.e., newer than the sequence mutations that define the tip branches – and frequent.

### The identification of positively selected genes

Many previous studies have implicated genes and their protein products in ongoing arms-races between pathogens and their hosts (84, 85). One way to approach this question is agnostic to function, which is to screen for genes with a history of positive selection. Ours is not the first attempt to detect selection in *X. fastidiosa* genomes. Previous studies have searched for selection by comparing levels of polymorphism or rates of synonymous and nonsynonymous mutations in the core genome using Tajima’s D and the McDonald-Kreitman test (24, 40). Other work has measured *ω* in core genes but without statistically testing for positive selection (83) or by applying the global test for ρο for > 1.0 (25). To our knowledge no other study of *X. fastidiosa* has tested for positive selection in accessory genes nor applied codon sites models. The set of positively selected *X. fastidiosa* genes represents candidate pathogenicity factors to mediate interactions with the environment, including plant host, insect vectors, or members of the microbial community.

To study positive selection, we estimated *ω*, or the ratio of nonsynonymous to synonymous mutations, for each core gene and for each accessory gene found in four or more isolates. In total, this exercise encompassed 5,135 genes: 1,257 core genes, 3,691 accessory genes, and 187 multicopy genes. We began by applying a global test that estimates *ω* over all sites and phylogenetic lineages. This approach can be overly conservative, because a significant test of *ω* > 1.0 requires that positive selection is very strong, acts across many sites in a gene, is present in most of the branches of the phylogeny, or all of the above. As expected, we found only a few genes (eight accessory genes in total) that were significant for positive selection with this test. Unfortunately, the annotations of 7 of 8 of these genes yielded few insights into their functions. To explore gene function further, we identified protein domains using the Conserved Domain Database. We found, for example, that the gene *group_7848* contains a VirB3 protein domain, which is part of the Type IV secretory pathway and is commonly associated with the membranes of the bacterial cell. The gene *cya* was also implicated using this test, which encodes adenylate cyclase and plays essential roles in regulation of cellular metabolism (86). Interestingly, the *cya* protein is involved in the cyclic AMP system, which is a global regulator in gram-negative bacteria and has been shown to modulate gene expression in pathogenic bacteria (87, 88).

The global test did allow, however, for two broad generalizations about patterns of selection in *X. fastidiosa*. First, as a group the core genes are under strong purifying selection with most (>90%) having *ω* estimates significantly < 1.0. Second, accessory genes generally have lower levels of purifying selection, as evidenced by a lower proportion (45%) of significant tests for *ω* < 1.0 and by much higher average 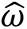 values (0.21973 vs. 0.51443; Figure 5A). The proportion of significant tests must be compared between genic sets with caution, because the smaller sample sizes (*n*=4 to 59) for accessory genes likely reduce statistical power relative to the minimum of 60 samples for all core genes, as do any differences in gene lengths. Nonetheless, the contrasting pattern of *ω* is consistent with the ideas that core genes have conserved biological functions and that accessory genes are more amenable to evolutionary change due to their nonessential, but still potentially biological relevant, cellular roles (89). Accessory genes may also experience higher variation in their selection dynamics because recombination affects them more than core genes (83).

Given few signals of positive selection with the global test, we turned to codon site models. To our surprise, the proportion of positively selected genes was similar for core genes (5.3%) and accessory genes (5.4%). The salient question is whether these genes give some clue to function. Of the 67 core genes with evidence for positive selection at the codon level, 40% were unannotated. We performed a functional analysis by grouping the protein coding sequences of these 67 core genes into COG categories to infer cellular functions. Excluding the category of unknown function, the largest category was ‘cell wall/membrane/envelope biogenesis,’ followed by the ‘amino acid metabolism and transport,’ ‘carbohydrate metabolism and transport,’ ‘translation,’ and ‘intracellular trafficking and secretion’ (Supplementary Figure 4A).

Of the 201 accessory genes with evidence for positive selection at the codon level, 82% were not annotated for function. The remaining set of 36 genes was enriched for GO categories related to protein secretion by the type IV secretion system (Supplementary Table 8). To better infer function, we performed a COG analysis and found that the largest categories (excluding the category of unknown function) were ‘intracellular trafficking and secretion,’ ‘replication, recombination and repair,’ and ‘secondary metabolites biosynthesis, transport and catabolism’ (Supplementary Figure 4B). Intriguingly, of this set of 201 genes, 50 overlapped with the set of 367 genes that had a gene-wide estimate of 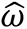 > 1. While these are especially strong candidates for having a history of positive selection, a disappointing 94% of them were unannotated for function. The three genes with annotations were: *cya*, *nagZ_2*, and *bacterial adaptive response A* (*barA)*. The gene *nagZ_2* encodes a beta-glucosidase that is important for biofilm formation in *Neisseria gonorrhoeae*, suggesting it could play a similar role in *X. fastidiosa*. It merits further functional analysis, since biofilms are important to the infection cycle (90). *barA* encodes a membrane associated histidine kinase that has a regulatory role in cell division, metabolism, and pili formation, and it has been implicated in regulating the virulence response of uropathogenic *E. coli* (91, 92). Finally, the multicopy genes also yielded evidence of positive selection, including *cdiA1*, which is part of the secretory contact-dependent growth inhibition (CDI) system that modulates biofilm formation in *Acinetobacter baumannii* (93).

As a final exercise, we cataloged the incidence of positive selection in a set of 35 genes that have been listed as virulence and pathogenicity factors in *X. fastidiosa* (13). Of the 35, we could identify 29 in our database based on the PD number annotation and reference sequence (http://www.microbesonline.org/operons/gnc183190.html; Table 1). We expected that this set of 29 genes would be enriched for evidence of positive selection relative to the genomic background, because these genes are putatively involved in arms-race interactions. The trend for these genes was in the expected direction, because 4 of 29 (= 13.9%) were significant vs. 301 of 5,135 (= 5.8%) in the rest of the genome. However, the difference in proportions was not significant (Fisher Exact Test, P = 0.1091). Nonetheless, this set of experimental genes is interesting. All four genes with evidence of positive selection encode proteins associated with the membrane of gram-negative bacteria and are involved in membrane transport or adhesin. Specifically, the genes *fimF, xadA* and *xatA* encode proteins involved in fimbrial adhesion, non-fimbrial adhesion, and biofilm formation, respectively, and the gene PD1311 encodes a protein involved in membrane transport (94–98). Because there is a resolved protein structure for fimF (99), we investigated the location of positively selected codons. Of the four positively selected codons (N80, D87, F137, and D142), one (D87) was in a flexible loop and a second (D142) comprised part of the second β-sheet of the protein (99). Together this suggests that changes in the amino acid sequence of *fimF* may be impacting its function

**Table 1:**
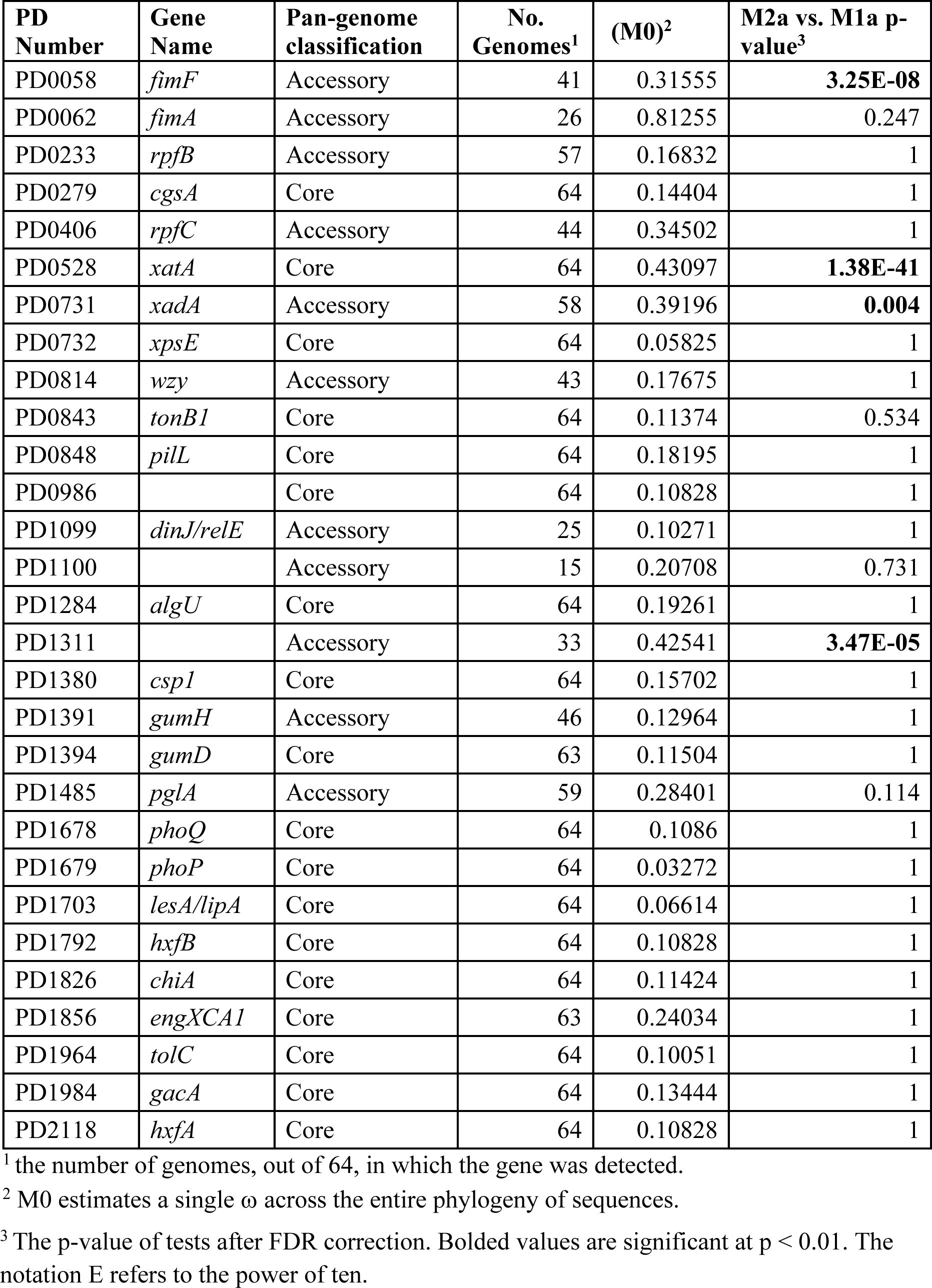
*Codeml* results for experimentally identified virulence and pathogenicity genes, as listed (13)

We must caution that positive selection analyses are subject to false positives, and they are also dependent on specific analysis features, like the sample set, the criteria for determining homology, and the sequence alignments. We have nonetheless found several genes with some evidence of positive selection that may also contribute to functions relevant to infection. We believe they represent suitable candidates for further functional analyses to elucidate their role in host-pathogen interactions and perhaps even host specificity.

## ACKNOWLEDGEMENTS

We thank three anonymous reviewers for comments. We also thank R. Gaut, INRA-CFBP, and the UC Irvine Genomics High-Throughput Facility for contributing to data generation. We also thank J.J. Emerson, A. Martiny, E. Solares, and C.I. Rodriguez for helpful input on methodology. T.N.B. was supported by the National Science Foundation Graduate Research Fellowship Program and the University of California Irvine President’s Dissertation Year Fellowship. This work was supported by National Science Foundation grant no. 1741627 to B.S.G. and California Department of Food and Agriculture agreement numbers 15-0218-SA and 18-0328-000-SA to M.C.R.

## REFERENCES

1. Furuya EY, Lowy FD. 2006. Antimicrobial-resistant bacteria in the community setting. 1. Nat Rev Microbiol 4:36–45.

2. Yacoubi BE, Brunings AM, Yuan Q, Shankar S, Gabriel DW. 2007. In Planta Horizontal Transfer of a Major Pathogenicity Effector Gene. Appl Environ Microbiol 73:1612–1621.

3. Juhas M. 2015. Horizontal gene transfer in human pathogens. Crit Rev Microbiol 41:101– 108.

4. Chen NWG, Serres-Giardi L, Ruh M, Briand M, Bonneau S, Darrasse A, Barbe V, Gagnevin L, Koebnik R, Jacques M-A. 2018. Horizontal gene transfer plays a major role in the pathological convergence of Xanthomonas lineages on common bean. BMC Genomics 19:606.

5. Welch RA, Burland V, Plunkett G, Redford P, Roesch P, Rasko D, Buckles EL, Liou S-R, Boutin A, Hackett J, Stroud D, Mayhew GF, Rose DJ, Zhou S, Schwartz DC, Perna NT, Mobley HLT, Donnenberg MS, Blattner FR. 2002. Extensive mosaic structure revealed by the complete genome sequence of uropathogenic Escherichia coli. Proc Natl Acad Sci 99:17020–17024.

6. Badet T, Croll D. 2020. The rise and fall of genes: origins and functions of plant pathogen pangenomes. Curr Opin Plant Biol 56:65–73.

7. Kim Y, Gu C, Kim HU, Lee SY. 2020. Current status of pan-genome analysis for pathogenic bacteria. Curr Opin Biotechnol 63:54–62.

8. Sicard A, Zeilinger AR, Vanhove M, Schartel TE, Beal DJ, Daugherty MP, Almeida RPP. 2018. Xylella fastidiosa: Insights into an Emerging Plant Pathogen. Annu Rev Phytopathol 56:181–202.

9. Burbank LP, Roper MC 2021. 2021. Microbe Profile: Xylella fastidiosa – a devastating agricultural pathogen with an endophytic lifestyle. Microbiology 167:001091.

10. Schuenzel EL, Scally M, Stouthamer R, Nunney L. 2005. A Multigene Phylogenetic Study of Clonal Diversity and Divergence in North American Strains of the Plant Pathogen Xylella fastidiosa. Appl Environ Microbiol 71:3832–3839.

11. Loconsole G, Saponari M, Boscia D, D’Attoma G, Morelli M, Martelli GP, Almeida RPP. 2016. Intercepted isolates of Xylella fastidiosa in Europe reveal novel genetic diversity. Eur J Plant Pathol 146:85–94.

12. Chatterjee S, Almeida RPP, Lindow S. 2008. Living in two Worlds: The Plant and Insect Lifestyles of Xylella fastidiosa. Annu Rev Phytopathol 46:243–271.

13. Rapicavoli J, Ingel B, Blanco-Ulate B, Cantu D, Roper C. 2018. Xylella fastidiosa: an examination of a re-emerging plant pathogen. Mol Plant Pathol 19:786–800.

14. Tumber K, Alston J, Fuller K. 2014. Pierce’s disease costs California $104 million per year. Calif Agric 68:20–29.

15. 2015. Assessing the returns to R&D on perennial crops: the costs and benefits of Pierce’s disease research in the California winegrape industry. Aust J Agric Resour Econ https://doi.org/10.22004/ag.econ.280230.

16. Koo H, Allan RN, Howlin RP, Stoodley P, Hall-Stoodley L. 2017. Targeting microbial biofilms: current and prospective therapeutic strategies. 12. Nat Rev Microbiol 15:740–755.

17. Castro C, DiSalvo B, Roper MC. 2021. Xylella fastidiosa: A reemerging plant pathogen that threatens crops globally. PLOS Pathog 17:e1009813.

18. Roper C, Lindow SE. 2016. CHAPTER 16: Xylella fastidiosa: Insights into the Lifestyle of a Xylem-Limited Bacterium, p. 307–320. In Caroline, R, Steven, EL (eds.), Virulence Mechanisms of Plant-Pathogenic Bacteria. The American Phytopathological Society.

19. Marcelletti S, Scortichini M. 2016. Genome-wide comparison and taxonomic relatedness of multiple Xylella fastidiosa strains reveal the occurrence of three subspecies and a new Xylella species. Arch Microbiol 198:803–812.

20. Nunney L, Vickerman DB, Bromley RE, Russell SA, Hartman JR, Morano LD, Stouthamer R. 2013. Recent Evolutionary Radiation and Host Plant Specialization in the Xylella fastidiosa Subspecies Native to the United States. Appl Environ Microbiol 79:2189–2200.

21. Almeida RPP, Purcell AH. 2003. Biological Traits of Xylella fastidiosa Strains from Grapes and Almonds. Appl Environ Microbiol 69:7447–7452.

22. Hernandez-Martinez R, Costa HS, Dumenyo CK, Cooksey DA. 2006. Differentiation of Strains of Xylella fastidiosa Infecting Grape, Almonds, and Oleander Using a Multiprimer PCR Assay. Plant Dis 90:1382–1388.

23. Almeida RPP, Nascimento FE, Chau J, Prado SS, Tsai C-W, Lopes SA, Lopes JRS. 2008. Genetic Structure and Biology of Xylella fastidiosa Strains Causing Disease in Citrus and Coffee in Brazil. Appl Environ Microbiol 74:3690–3701.

24. Castillo AI, Bojanini I, Chen H, Kandel PP, De La Fuente L, Almeida RPP. 2021. Allopatric Plant Pathogen Population Divergence following Disease Emergence. Appl Environ Microbiol 87:e02095–20.

25. Vanhove M, Sicard A, Ezennia J, Leviten N, Almeida RPP. 2020. Population structure and adaptation of a bacterial pathogen in California grapevines. Environ Microbiol 22:2625– 2638.

26. Uceda-Campos G, Feitosa-Junior OR, Santiago CRN, Pierry PM, Zaini PA, de Santana WO, Martins-Junior J, Barbosa D, Digiampietri LA, Setubal JC, da Silva AM. 2022. Comparative Genomics of Xylella fastidiosa Explores Candidate Host-Specificity Determinants and Expands the Known Repertoire of Mobile Genetic Elements and Immunity Systems. 5. Microorganisms 10:914.

27. Kahn AK, Almeida RPP. 2022. Phylogenetics of Historical Host Switches in a Bacterial Plant Pathogen. Appl Environ Microbiol 88:e02356–21.

28. Daugherty MD, Malik HS. 2012. Rules of Engagement: Molecular Insights from Host-Virus Arms Races. Annu Rev Genet 46:677–700.

29. Aleru O, Barber MF. 2020. Battlefronts of evolutionary conflict between bacteria and animal hosts. PLOS Pathog 16:e1008797.

30. Mitchell PS, Patzina C, Emerman M, Haller O, Malik HS, Kochs G. 2012. Evolution-Guided Identification of Antiviral Specificity Determinants in the Broadly Acting Interferon-Induced Innate Immunity Factor MxA. Cell Host Microbe 12:598–604.

31. Ng M, Ndungo E, Kaczmarek ME, Herbert AS, Binger T, Kuehne AI, Jangra RK, Hawkins JA, Gifford RJ, Biswas R, Demogines A, James RM, Yu M, Brummelkamp TR, Drosten C, Wang L-F, Kuhn JH, Müller MA, Dye JM, Sawyer SL, Chandran K. 2015. Filovirus receptor NPC1 contributes to species-specific patterns of ebolavirus susceptibility in bats. eLife 4:e11785.

32. Daugherty MD, Schaller AM, Geballe AP, Malik HS. 2016. Evolution-guided functional analyses reveal diverse antiviral specificities encoded by IFIT1 genes in mammals. eLife. eLife Sciences Publications Limited. https://elifesciences.org/articles/14228. Retrieved 2 April 2022.

33. Andrews S. 2010. FastQC: A Quality Control Tool for High Throughput Sequence Data. http://www.bioinformatics.babraham.ac.uk/projects/fastqc/.

34. Bolger AM, Lohse M, Usadel B. 2014. Trimmomatic: a flexible trimmer for Illumina sequence data. Bioinformatics 30:2114–2120.

35. Koren S, Walenz BP, Berlin K, Miller JR, Bergman NH, Phillippy AM. 2017. Canu: scalable and accurate long-read assembly via adaptive k-mer weighting and repeat separation. Genome Res gr.215087.116.

36. Wick RR, Judd LM, Gorrie CL, Holt KE. 2017. Unicycler: Resolving bacterial genome assemblies from short and long sequencing reads. PLOS Comput Biol 13:e1005595.

37. Gurevich A, Saveliev V, Vyahhi N, Tesler G. 2013. QUAST: quality assessment tool for genome assemblies. Bioinforma Oxf Engl 29:1072–1075.

38. Chase AB, Gomez-Lunar Z, Lopez AE, Li J, Allison SD, Martiny AC, Martiny JBH. 2018. Emergence of soil bacterial ecotypes along a climate gradient. Environ Microbiol 20:4112– 4126.

39. Shen W, Le S, Li Y, Hu F. 2016. SeqKit: A Cross-Platform and Ultrafast Toolkit for FASTA/Q File Manipulation. PLOS ONE 11:e0163962.

40. Castillo AI, Chacón-Díaz C, Rodríguez-Murillo N, Coletta-Filho HD, Almeida RPP. 2020. Impacts of local population history and ecology on the evolution of a globally dispersed pathogen. BMC Genomics 21:369.

41. Bankevich A, Nurk S, Antipov D, Gurevich AA, Dvorkin M, Kulikov AS, Lesin VM, Nikolenko SI, Pham S, Prjibelski AD, Pyshkin AV, Sirotkin AV, Vyahhi N, Tesler G, Alekseyev MA, Pevzner PA. 2012. SPAdes: A New Genome Assembly Algorithm and Its Applications to Single-Cell Sequencing. J Comput Biol 19:455–477.

42. Seemann T. 2014. Prokka: rapid prokaryotic genome annotation. Bioinformatics 30:2068– 2069.

43. Page AJ, Cummins CA, Hunt M, Wong VK, Reuter S, Holden MTG, Fookes M, Falush D, Keane JA, Parkhill J. 2015. Roary: rapid large-scale prokaryote pan genome analysis. Bioinformatics 31:3691–3693.

44. Castresana J. 2000. Selection of Conserved Blocks from Multiple Alignments for Their Use in Phylogenetic Analysis. Mol Biol Evol 17:540–552.

45. Katoh K, Misawa K, Kuma K, Miyata T. 2002. MAFFT: a novel method for rapid multiple sequence alignment based on fast Fourier transform. Nucleic Acids Res 30:3059–3066.

46. Katoh K, Standley DM. 2013. MAFFT Multiple Sequence Alignment Software Version 7: Improvements in Performance and Usability. Mol Biol Evol 30:772–780.

47. Stamatakis A. 2014. RAxML version 8: a tool for phylogenetic analysis and post-analysis of large phylogenies. Bioinformatics 30:1312–1313.

48. Boc A, Diallo AB, Makarenkov V. 2012. T-REX: a web server for inferring, validating and visualizing phylogenetic trees and networks. Nucleic Acids Res 40:W573–W579.

49. Parks DH, Imelfort M, Skennerton CT, Hugenholtz P, Tyson GW. 2015. CheckM: assessing the quality of microbial genomes recovered from isolates, single cells, and metagenomes. Genome Res 25:1043–1055.

50. Rodriguez CI, Martiny JBH. 2020. Evolutionary relationships among bifidobacteria and their hosts and environments. BMC Genomics 21:26.

51. Madeira F, Park Y mi, Lee J, Buso N, Gur T, Madhusoodanan N, Basutkar P, Tivey ARN, Potter SC, Finn RD, Lopez R. 2019. The EMBL-EBI search and sequence analysis tools APIs in 2019. Nucleic Acids Res 47:W636–W641.

52. Huerta-Cepas J, Forslund K, Coelho LP, Szklarczyk D, Jensen LJ, von Mering C, Bork P. 2017. Fast Genome-Wide Functional Annotation through Orthology Assignment by eggNOG-Mapper. Mol Biol Evol 34:2115–2122.

53. Huerta-Cepas J, Szklarczyk D, Heller D, Hernández-Plaza A, Forslund SK, Cook H, Mende DR, Letunic I, Rattei T, Jensen LJ, von Mering C, Bork P. 2019. eggNOG 5.0: a hierarchical, functionally and phylogenetically annotated orthology resource based on 5090 organisms and 2502 viruses. Nucleic Acids Res 47:D309–D314.

54. Ashburner M, Ball CA, Blake JA, Botstein D, Butler H, Cherry JM, Davis AP, Dolinski K, Dwight SS, Eppig JT, Harris MA, Hill DP, Issel-Tarver L, Kasarskis A, Lewis S, Matese JC, Richardson JE, Ringwald M, Rubin GM, Sherlock G. 2000. Gene Ontology: tool for the unification of biology. Nat Genet 25:25–29.

55. Nguyen L-T, Schmidt HA, von Haeseler A, Minh BQ. 2015. IQ-TREE: A Fast and Effective Stochastic Algorithm for Estimating Maximum-Likelihood Phylogenies. Mol Biol Evol 32:268–274.

56. Kalyaanamoorthy S, Minh BQ, Wong TKF, von Haeseler A, Jermiin LS. 2017. ModelFinder: fast model selection for accurate phylogenetic estimates. 6. Nat Methods 14:587–589.

57. Paradis E, Schliep K. 2019. ape 5.0: an environment for modern phylogenetics and evolutionary analyses in R. Bioinformatics 35:526–528.

58. R Core Team. 2019. R: A language and environment for statistical computing. R Foundation for Statistical Computing. https://www.R-project.org/.

59. Oksanen J, Guillaume Blanchet F, Friendly M, Kindt R, Legendre P, McGlinn D, Minchin PR, O’Hara RB, Simpson GL, Solymos P, Stevens MHH, Szoecs E, Wagner H. 2020. vegan: Community Ecology Package.

60. Kung SH, Almeida RPP. 2011. Natural Competence and Recombination in the Plant Pathogen Xylella fastidiosa. Appl Environ Microbiol 77:5278–5284.

61. Croucher NJ, Page AJ, Connor TR, Delaney AJ, Keane JA, Bentley SD, Parkhill J, Harris SR. 2015. Rapid phylogenetic analysis of large samples of recombinant bacterial whole genome sequences using Gubbins. Nucleic Acids Res 43:e15.

62. Shikov AE, Malovichko YV, Nizhnikov AA, Antonets KS. 2022. Current Methods for Recombination Detection in Bacteria. 11. Int J Mol Sci 23:6257.

63. Revell LJ. 2012. phytools: an R package for phylogenetic comparative biology (and other things). Methods Ecol Evol 3:217–223.

64. Mateo-Estrada V, Graña-Miraglia L, López-Leal G, Castillo-Ramírez S. 2019. Phylogenomics Reveals Clear Cases of Misclassification and Genus-Wide Phylogenetic Markers for Acinetobacter. Genome Biol Evol 11:2531–2541.

65. Bouckaert RR. 2010. DensiTree: making sense of sets of phylogenetic trees. Bioinformatics 26:1372–1373.

66. Löytynoja A. 2014. Phylogeny-aware alignment with PRANK, p. 155–170. *In* Russell, DJ (ed.), Multiple Sequence Alignment Methods. Humana Press, Totowa, NJ.

67. Schliep KP. 2011. phangorn: phylogenetic analysis in R. Bioinformatics 27:592–593.

68. Cohen O, Ashkenazy H, Belinky F, Huchon D, Pupko T. 2010. GLOOME: gain loss mapping engine. Bioinformatics 26:2914–2915.

69. Avram O, Rapoport D, Portugez S, Pupko T. 2019. M1CR0B1AL1Z3R—a user-friendly web server for the analysis of large-scale microbial genomics data. Nucleic Acids Res 47:W88–W92.

70. Yang Z. 1997. PAML: a program package for phylogenetic analysis by maximum likelihood. Bioinformatics 13:555–556.

71. Yang Z. 2007. PAML 4: Phylogenetic Analysis by Maximum Likelihood. Mol Biol Evol 24:1586–1591.

72. Denancé N, Briand M, Gaborieau R, Gaillard S, Jacques M-A. 2019. Identification of genetic relationships and subspecies signatures in Xylella fastidiosa. BMC Genomics 20:239.

73. Bolotin E, Hershberg R. 2015. Gene Loss Dominates As a Source of Genetic Variation within Clonal Pathogenic Bacterial Species. Genome Biol Evol 7:2173–2187.

74. Iranzo J, Wolf YI, Koonin EV, Sela I. 2019. Gene gain and loss push prokaryotes beyond the homologous recombination barrier and accelerate genome sequence divergence. Nat Commun 10:5376.

75. Firrao G, Scortichini M, Pagliari L. 2021. Orthology-Based Estimate of the Contribution of Horizontal Gene Transfer from Distantly Related Bacteria to the Intraspecific Diversity and Differentiation of Xylella fastidiosa. 1. Pathogens 10:46.

76. Graña-Miraglia L, Lozano LF, Velázquez C, Volkow-Fernández P, Pérez-Oseguera Á, Cevallos MA, Castillo-Ramírez S. 2017. Rapid Gene Turnover as a Significant Source of Genetic Variation in a Recently Seeded Population of a Healthcare-Associated Pathogen. Front Microbiol 8.

77. Sironi M, Cagliani R, Forni D, Clerici M. 2015. Evolutionary insights into host–pathogen interactions from mammalian sequence data. 4. Nat Rev Genet 16:224–236.

78. Giampetruzzi A, Saponari M, Loconsole G, Boscia D, Savino VN, Almeida RPP, Zicca S, Landa BB, Chacón-Diaz C, Saldarelli P. 2017. Genome-Wide Analysis Provides Evidence on the Genetic Relatedness of the Emergent Xylella fastidiosa Genotype in Italy to Isolates from Central America. Phytopathology® 107:816–827.

79. Tettelin H, Riley D, Cattuto C, Medini D. 2008. Comparative genomics: the bacterial pan-genome. Curr Opin Microbiol 11:472–477.

80. Yuan X, Morano L, Bromley R, Spring-Pearson S, Stouthamer R, Nunney L. 2010. Multilocus sequence typing of Xylella fastidiosa causing Pierce’s disease and oleander leaf scorch in the United States. Phytopathology 100:601–611.

81. Hurles M. 2004. Gene Duplication: The Genomic Trade in Spare Parts. PLOS Biol 2:e206.

82. Arnold BJ, Huang I-T, Hanage WP. 2022. Horizontal gene transfer and adaptive evolution in bacteria. 4. Nat Rev Microbiol 20:206–218.

83. Castillo AI, Almeida RPP. 2021. Evidence of gene nucleotide composition favoring replication and growth in a fastidious plant pathogen. G3 GenesGenomesGenetics 11:jkab076.

84. Anderson JP, Gleason CA, Foley RC, Thrall PH, Burdon JB, Singh KB, Anderson JP, Gleason CA, Foley RC, Thrall PH, Burdon JB, Singh KB. 2010. Plants versus pathogens: an evolutionary arms race. Funct Plant Biol 37:499–512.

85. Schulte RD, Makus C, Hasert B, Michiels NK, Schulenburg H. 2010. Multiple reciprocal adaptations and rapid genetic change upon experimental coevolution of an animal host and its microbial parasite. Proc Natl Acad Sci 107:7359–7364.

86. Danchin A, Guiso N, Roy A, Ullmann A. 1984. Identification of the Escherichia coli cya gene product as authentic adenylate cyclase. J Mol Biol 175:403–408.

87. Smith RS, Wolfgang MC, Lory S. 2004. An Adenylate Cyclase-Controlled Signaling Network Regulates Pseudomonas aeruginosa Virulence in a Mouse Model of Acute Pneumonia. Infect Immun 72:1677–1684.

88. Kim YR, Kim SY, Kim CM, Lee SE, Rhee JH. 2005. Essential role of an adenylate cyclase in regulating Vibrio vulnificus virulence. FEMS Microbiol Lett 243:497–503.

89. Horesh G, Taylor-Brown A, McGimpsey S, Lassalle F, Corander J, Heinz E, Thomson NR. 2021. Different evolutionary trends form the twilight zone of the bacterial pan-genome. Microb Genomics 7:000670.

90. Bhoopalan SV, Piekarowicz A, Lenz JD, Dillard JP, Stein DC. 2016. nagZ Triggers Gonococcal Biofilm Disassembly. 1. Sci Rep 6:22372.

91. Palaniyandi S, Mitra A, Herren CD, Lockatell CV, Johnson DE, Zhu X, Mukhopadhyay S. 2012. BarA-UvrY Two-Component System Regulates Virulence of Uropathogenic E. coli CFT073. PLOS ONE 7:e31348.

92. Sahu SN, Acharya S, Tuminaro H, Patel I, Dudley K, LeClerc JE, Cebula TA, Mukhopadhyay S. 2003. The bacterial adaptive response gene, barA, encodes a novel conserved histidine kinase regulatory switch for adaptation and modulation of metabolism in Escherichia coli. Mol Cell Biochem 253:167–177.

93. Roussin M, Rabarioelina S, Cluzeau L, Cayron J, Lesterlin C, Salcedo SP, Bigot S. 2019. Identification of a Contact-Dependent Growth Inhibition (CDI) System That Reduces Biofilm Formation and Host Cell Adhesion of Acinetobacter baumannii DSM30011 Strain. Front Microbiol 10:2450.

94. Ma Rodriguez A, Olano C, Vilches C, Méndez C, Salas JA. 1993. Streptomyces antibioticus contains at least three oleandomycin-resistance determinants, one of which shows similarity with proteins of the ABC-transporter superfamily. Mol Microbiol 8:571– 582.

95. Sun QH, Hu J, Huang GX, Ge C, Fang RX, He CZ. 2005. Type-II secretion pathway structural gene xpsE, xylanase- and cellulase secretion and virulence in Xanthomonas oryzae pv. oryzae. Plant Pathol 54:15–21.

96. Abbas A, Adams C, Scully N, Glennon J, O’Gara F. 2007. A role for TonB1 in biofilm formation and quorum sensing in Pseudomonas aeruginosa. FEMS Microbiol Lett 274:269– 278.

97. Das A, Rangaraj N, Sonti RV. 2009. Multiple Adhesin-Like Functions of *Xanthomonas oryzae* pv. *oryzae* Are Involved in Promoting Leaf Attachment, Entry, and Virulence on Rice. Mol Plant-Microbe Interactions® 22:73–85.

98. Zeiner SA, Dwyer BE, Clegg S. 2012. FimA, FimF, and FimH Are Necessary for Assembly of Type 1 Fimbriae on Salmonella enterica Serovar Typhimurium. Infect Immun 80:3289–3296.

99. Gossert AD, Bettendorff P, Puorger C, Vetsch M, Herrmann T, Glockshuber R, Wüthrich K. 2008. NMR Structure of the Escherichia coli Type 1 Pilus Subunit FimF and Its Interactions with Other Pilus Subunits. J Mol Biol 375:752–763.

