## Supplementary Materials for "Using genomes and evolutionary analyses to screen for host-specificity and positive selection in the plant pathogen *Xylella fastidiosa*"

**Supplementary Table 5:** GO analysis of core genes

| GO biological process complete | REFLIST | upload | expected | over/under | fold Enrichment | P-value |
|---|---|---|---|---|---|---|
| pyrimidine-containing compound biosynthetic process (GO:0072528) | 20 | 16 | 2.99 | + | 5.35 | 1.91E-03 |
| translation (GO:0006412) | 105 | 73 | 15.71 | + | 4.65 | 8.14E-19 |
| aromatic amino acid family biosynthetic process (GO:0009073) | 22 | 15 | 3.29 | + | 4.56 | 1.51E-02 |
| regulation of developmental process (GO:0050793) | 28 | 19 | 4.19 | + | 4.54 | 1.32E-03 |
| peptide biosynthetic process (GO:0043043) | 113 | 76 | 16.91 | + | 4.5 | 4.72E-19 |
| regulation of cell shape (GO:0008360) | 27 | 18 | 4.04 | + | 4.46 | 2.93E-03 |
| regulation of cell morphogenesis (GO:0022604) | 27 | 18 | 4.04 | + | 4.46 | 2.93E-03 |
| regulation of anatomical structure morphogenesis (GO:0022603) | 27 | 18 | 4.04 | + | 4.46 | 2.93E-03 |
| ribonucleoside monophosphate metabolic process (GO:0009161) | 29 | 19 | 4.34 | + | 4.38 | 1.93E-03 |
| amide biosynthetic process (GO:0043604) | 140 | 91 | 20.95 | + | 4.34 | 1.53E-22 |
| nucleoside monophosphate biosynthetic process (GO:0009124) | 31 | 20 | 4.64 | + | 4.31 | 1.26E-03 |
| peptide metabolic process (GO:0006518) | 133 | 85 | 19.9 | + | 4.27 | 1.63E-20 |
| nucleoside monophosphate metabolic process (GO:0009123) | 36 | 23 | 5.39 | + | 4.27 | 2.46E-04 |
| RNA methylation (GO:0001510) | 22 | 14 | 3.29 | + | 4.25 | 4.84E-02 |
| ribonucleoside monophosphate biosynthetic process (GO:0009156) | 27 | 17 | 4.04 | + | 4.21 | 9.32E-03 |
| pyrimidine-containing compound metabolic process (GO:0072527) | 27 | 17 | 4.04 | + | 4.21 | 9.32E-03 |
| cell cycle (GO:0007049) | 42 | 26 | 6.28 | + | 4.14 | 6.80E-05 |
| cellular amide metabolic process (GO:0043603) | 178 | 104 | 26.63 | + | 3.91 | 2.08E-23 |
| ribonucleotide biosynthetic process (GO:0009260) | 52 | 30 | 7.78 | + | 3.86 | 2.33E-05 |
| ribose phosphate biosynthetic process (GO:0046390) | 53 | 30 | 7.93 | + | 3.78 | 3.21E-05 |
| ribonucleotide metabolic process (GO:0009259) | 78 | 44 | 11.67 | + | 3.77 | 2.47E-08 |
| glycosaminoglycan biosynthetic process (GO:0006024) | 32 | 18 | 4.79 | + | 3.76 | 1.64E-02 |
| aminoglycan biosynthetic process (GO:0006023) | 32 | 18 | 4.79 | + | 3.76 | 1.64E-02 |
| peptidoglycan biosynthetic process (GO:0009252) | 32 | 18 | 4.79 | + | 3.76 | 1.64E-02 |
| ribosome biogenesis (GO:0042254) | 47 | 26 | 7.03 | + | 3.7 | 3.50E-04 |
| ribonucleoprotein complex biogenesis (GO:0022613) | 47 | 26 | 7.03 | + | 3.7 | 3.50E-04 |
| ribose phosphate metabolic process (GO:0019693) | 82 | 45 | 12.27 | + | 3.67 | 2.94E-08 |
| purine ribonucleotide biosynthetic process (GO:0009152) | 42 | 23 | 6.28 | + | 3.66 | 1.79E-03 |

| | | | | | | |
|---|---|---|---|---|---|---|
| monocarboxylic acid biosynthetic process (GO:0072330) | 42 | 23 | 6.28 | + | 3.66 | 1.79E-03 |
| cellular component macromolecule biosynthetic process (GO:0070589) | 33 | 18 | 4.94 | + | 3.65 | 2.23E-02 |
| purine ribonucleotide metabolic process (GO:0009150) | 66 | 36 | 9.87 | + | 3.65 | 3.05E-06 |
| cell wall macromolecule biosynthetic process (GO:0044038) | 33 | 18 | 4.94 | + | 3.65 | 2.23E-02 |
| cellular macromolecule biosynthetic process (GO:0034645) | 178 | 97 | 26.63 | + | 3.64 | 6.22E-20 |
| gene expression (GO:0010467) | 244 | 132 | 36.5 | + | 3.62 | 3.41E-28 |
| tRNA metabolic process (GO:0006399) | 65 | 35 | 9.72 | + | 3.6 | 6.45E-06 |
| organonitrogen compound biosynthetic process (GO:1901566) | 406 | 218 | 60.74 | + | 3.59 | 4.74E-51 |
| sulfur compound biosynthetic process (GO:0044272) | 56 | 30 | 8.38 | + | 3.58 | 8.05E-05 |
| nucleotide biosynthetic process (GO:0009165) | 71 | 38 | 10.62 | + | 3.58 | 1.69E-06 |
| peptidoglycan-based cell wall biogenesis (GO:0009273) | 36 | 19 | 5.39 | + | 3.53 | 1.92E-02 |
| nucleoside phosphate biosynthetic process (GO:1901293) | 72 | 38 | 10.77 | + | 3.53 | 2.27E-06 |
| tRNA processing (GO:0008033) | 40 | 21 | 5.98 | + | 3.51 | 7.90E-03 |
| purine nucleotide biosynthetic process (GO:0006164) | 44 | 23 | 6.58 | + | 3.49 | 3.25E-03 |
| vitamin biosynthetic process (GO:0009110) | 46 | 24 | 6.88 | + | 3.49 | 2.08E-03 |
| water-soluble vitamin biosynthetic process (GO:0042364) | 46 | 24 | 6.88 | + | 3.49 | 2.08E-03 |
| purine nucleotide metabolic process (GO:0006163) | 69 | 36 | 10.32 | + | 3.49 | 7.43E-06 |
| cellular amino acid biosynthetic process (GO:0008652) | 100 | 52 | 14.96 | + | 3.48 | 4.04E-09 |
| cell division (GO:0051301) | 54 | 28 | 8.08 | + | 3.47 | 3.51E-04 |
| cellular nitrogen compound biosynthetic process (GO:0044271) | 365 | 189 | 54.61 | + | 3.46 | 5.09E-41 |
| cell wall biogenesis (GO:0042546) | 37 | 19 | 5.54 | + | 3.43 | 2.56E-02 |
| phospholipid biosynthetic process (GO:0008654) | 45 | 23 | 6.73 | + | 3.42 | 4.32E-03 |
| phospholipid metabolic process (GO:0006644) | 45 | 23 | 6.73 | + | 3.42 | 4.32E-03 |
| organophosphate biosynthetic process (GO:0090407) | 135 | 69 | 20.2 | + | 3.42 | 2.08E-12 |
| purine-containing compound biosynthetic process (GO:0072522) | 47 | 24 | 7.03 | + | 3.41 | 2.77E-03 |
| water-soluble vitamin metabolic process (GO:0006767) | 49 | 25 | 7.33 | + | 3.41 | 1.77E-03 |
| vitamin metabolic process (GO:0006766) | 49 | 25 | 7.33 | + | 3.41 | 1.77E-03 |
| ncRNA metabolic process (GO:0034660) | 98 | 50 | 14.66 | + | 3.41 | 1.78E-08 |
| alpha-amino acid biosynthetic process (GO:1901607) | 85 | 43 | 12.72 | + | 3.38 | 5.63E-07 |

| | | | | | | |
|---|---|---|---|---|---|---|
| carboxylic acid biosynthetic process (GO:0046394) | 157 | 79 | 23.49 | + | 3.36 | 3.26E-14 |
| organic acid biosynthetic process (GO:0016053) | 159 | 79 | 23.79 | + | 3.32 | 5.71E-14 |
| RNA processing (GO:0006396) | 77 | 38 | 11.52 | + | 3.3 | 9.26E-06 |
| ncRNA processing (GO:0034470) | 73 | 36 | 10.92 | + | 3.3 | 2.27E-05 |
| external encapsulating structure organization (GO:0045229) | 41 | 20 | 6.13 | + | 3.26 | 2.84E-02 |
| nucleotide metabolic process (GO:0009117) | 107 | 52 | 16.01 | + | 3.25 | 2.83E-08 |
| small molecule biosynthetic process (GO:0044283) | 225 | 109 | 33.66 | + | 3.24 | 1.31E-19 |
| sulfur compound metabolic process (GO:0006790) | 93 | 45 | 13.91 | + | 3.23 | 6.77E-07 |
| carbohydrate derivative biosynthetic process (GO:1901137) | 152 | 73 | 22.74 | + | 3.21 | 3.79E-12 |
| dicarboxylic acid metabolic process (GO:0043648) | 50 | 24 | 7.48 | + | 3.21 | 6.23E-03 |
| nucleoside phosphate metabolic process (GO:0006753) | 111 | 53 | 16.61 | + | 3.19 | 3.04E-08 |
| purine-containing compound metabolic process (GO:0072521) | 78 | 37 | 11.67 | + | 3.17 | 3.20E-05 |
| cellular biosynthetic process (GO:0044249) | 616 | 292 | 92.16 | + | 3.17 | 1.39E-64 |
| organic substance biosynthetic process (GO:1901576) | 625 | 294 | 93.51 | + | 3.14 | 1.42E-64 |
| organophosphate metabolic process (GO:0019637) | 185 | 87 | 27.68 | + | 3.14 | 2.42E-14 |
| macromolecule biosynthetic process (GO:0009059) | 251 | 118 | 37.55 | + | 3.14 | 1.49E-20 |
| aromatic compound biosynthetic process (GO:0019438) | 235 | 109 | 35.16 | + | 3.1 | 2.01E-18 |
| organic cyclic compound biosynthetic process (GO:1901362) | 266 | 122 | 39.8 | + | 3.07 | 1.01E-20 |
| biosynthetic process (GO:0009058) | 650 | 296 | 97.25 | + | 3.04 | 1.50E-62 |
| heterocycle biosynthetic process (GO:0018130) | 251 | 113 | 37.55 | + | 3.01 | 2.33E-18 |
| RNA modification (GO:0009451) | 56 | 25 | 8.38 | + | 2.98 | 1.74E-02 |
| RNA metabolic process (GO:0016070) | 170 | 75 | 25.43 | + | 2.95 | 6.31E-11 |
| nucleobase-containing small molecule metabolic process (GO:0055086) | 147 | 64 | 21.99 | + | 2.91 | 7.92E-09 |
| cellular component biogenesis (GO:0044085) | 148 | 64 | 22.14 | + | 2.89 | 9.35E-09 |
| DNA repair (GO:0006281) | 72 | 31 | 10.77 | + | 2.88 | 2.25E-03 |
| nucleobase-containing compound biosynthetic process (GO:0034654) | 165 | 71 | 24.69 | + | 2.88 | 6.94E-10 |
| nucleic acid phosphodiester bond hydrolysis (GO:0090305) | 56 | 24 | 8.38 | + | 2.86 | 3.43E-02 |
| regulation of biological quality (GO:0065008) | 73 | 31 | 10.92 | + | 2.84 | 2.71E-03 |
| carbohydrate derivative metabolic process (GO:1901135) | 231 | 98 | 34.56 | + | 2.84 | 3.48E-14 |
| lipid biosynthetic process (GO:0008610) | 95 | 40 | 14.21 | + | 2.81 | 1.80E-04 |
| cellular amino acid metabolic process (GO:0006520) | 190 | 80 | 28.43 | + | 2.81 | 5.80E-11 |

| | | | | | | |
|---|---|---|---|---|---|---|
| cellular nitrogen compound metabolic process (GO:0034641) | 703 | 295 | 105.18 | + | 2.8 | 1.13E-55 |
| alpha-amino acid metabolic process (GO:1901605) | 138 | 57 | 20.65 | + | 2.76 | 6.99E-07 |
| cellular component organization or biogenesis (GO:0071840) | 197 | 81 | 29.47 | + | 2.75 | 1.33E-10 |
| carboxylic acid metabolic process (GO:0019752) | 327 | 134 | 48.92 | + | 2.74 | 2.14E-19 |
| oxoacid metabolic process (GO:0043436) | 334 | 136 | 49.97 | + | 2.72 | 1.32E-19 |
| small molecule metabolic process (GO:0044281) | 507 | 203 | 75.85 | + | 2.68 | 2.22E-31 |
| cellular response to DNA damage stimulus (GO:0006974) | 80 | 32 | 11.97 | + | 2.67 | 6.09E-03 |
| organic acid metabolic process (GO:0006082) | 341 | 136 | 51.02 | + | 2.67 | 6.21E-19 |
| cellular component organization (GO:0016043) | 152 | 60 | 22.74 | + | 2.64 | 9.23E-07 |
| monocarboxylic acid metabolic process (GO:0032787) | 123 | 48 | 18.4 | + | 2.61 | 5.03E-05 |
| cellular response to stress (GO:0033554) | 107 | 41 | 16.01 | + | 2.56 | 7.27E-04 |
| organic cyclic compound metabolic process (GO:1901360) | 614 | 233 | 91.86 | + | 2.54 | 2.41E-34 |
| heterocycle metabolic process (GO:0046483) | 589 | 223 | 88.12 | + | 2.53 | 3.20E-32 |
| cellular aromatic compound metabolic process (GO:0006725) | 587 | 220 | 87.82 | + | 2.51 | 4.84E-31 |
| cellular lipid metabolic process (GO:0044255) | 127 | 47 | 19 | + | 2.47 | 2.53E-04 |
| cellular protein metabolic process (GO:0044267) | 255 | 93 | 38.15 | + | 2.44 | 5.67E-10 |
| nucleobase-containing compound metabolic process (GO:0006139) | 495 | 179 | 74.06 | + | 2.42 | 2.37E-22 |
| organonitrogen compound metabolic process (GO:1901564) | 781 | 280 | 116.85 | + | 2.4 | 4.72E-40 |
| cellular metabolic process (GO:0044237) | 1323 | 455 | 197.93 | + | 2.3 | 1.26E-83 |
| nitrogen compound metabolic process (GO:0006807) | 1124 | 384 | 168.16 | + | 2.28 | 2.39E-60 |
| response to stress (GO:0006950) | 143 | 48 | 21.39 | + | 2.24 | 2.57E-03 |
| nucleic acid metabolic process (GO:0090304) | 354 | 118 | 52.96 | + | 2.23 | 3.85E-11 |
| phosphorus metabolic process (GO:0006793) | 328 | 107 | 49.07 | + | 2.18 | 2.48E-09 |
| cellular macromolecule metabolic process (GO:0044260) | 567 | 184 | 84.83 | + | 2.17 | 1.59E-18 |
| phosphate-containing compound metabolic process (GO:0006796) | 320 | 103 | 47.88 | + | 2.15 | 1.26E-08 |
| lipid metabolic process (GO:0006629) | 150 | 48 | 22.44 | + | 2.14 | 6.63E-03 |
| organic substance metabolic process (GO:0071704) | 1435 | 449 | 214.69 | + | 2.09 | 7.90E-69 |
| primary metabolic process (GO:0044238) | 1277 | 394 | 191.05 | + | 2.06 | 6.86E-52 |
| protein metabolic process (GO:0019538) | 383 | 116 | 57.3 | + | 2.02 | 1.81E-08 |
| metabolic process (GO:0008152) | 1590 | 477 | 237.88 | + | 2.01 | 3.77E-72 |
| macromolecule metabolic process (GO:0043170) | 821 | 239 | 122.83 | + | 1.95 | 2.28E-20 |
| cellular process (GO:0009987) | 1918 | 543 | 286.95 | + | 1.89 | 2.91E-90 |
| biological_process (GO:0008150) | 2336 | 582 | 349.49 | + | 1.67 | 3.82E-88 |
| Unclassified (UNCLASSIFIED) | 1768 | 32 | 264.51 | - | 0.12 | 0.00E+00 |

**Supplementary Table 6:** GO analysis of accessory genes

| GO biological process complete | REFLIST | upload | expected | over/under | fold Enrichment | P-value |
|---|---|---|---|---|---|---|
| glutamine family amino acid biosynthetic process (GO:0009084) | 17 | 8 | 0.94 | + | 8.51 | 1.46E-02 |
| glutamine metabolic process (GO:0006541) | 17 | 8 | 0.94 | + | 8.51 | 1.46E-02 |
| tRNA aminoacylation for protein translation (GO:0006418) | 25 | 10 | 1.38 | + | 7.23 | 4.31E-03 |
| tRNA aminoacylation (GO:0043039) | 25 | 10 | 1.38 | + | 7.23 | 4.31E-03 |
| amino acid activation (GO:0043038) | 26 | 10 | 1.44 | + | 6.95 | 5.69E-03 |
| chromosome organization (GO:0051276) | 27 | 9 | 1.49 | + | 6.03 | 4.03E-02 |
| glutamine family amino acid metabolic process (GO:0009064) | 37 | 11 | 2.05 | + | 5.37 | 1.51E-02 |
| purine nucleotide biosynthetic process (GO:0006164) | 44 | 12 | 2.43 | + | 4.93 | 1.34E-02 |
| tRNA metabolic process (GO:0006399) | 65 | 17 | 3.6 | + | 4.73 | 4.18E-04 |
| purine-containing compound biosynthetic process (GO:0072522) | 47 | 12 | 2.6 | + | 4.62 | 2.33E-02 |
| ribose phosphate biosynthetic process (GO:0046390) | 53 | 13 | 2.93 | + | 4.43 | 1.59E-02 |
| ncRNA metabolic process (GO:0034660) | 98 | 24 | 5.42 | + | 4.43 | 5.69E-06 |
| nucleotide biosynthetic process (GO:0009165) | 71 | 15 | 3.93 | + | 3.82 | 1.76E-02 |
| cellular amino acid biosynthetic process (GO:0008652) | 100 | 21 | 5.53 | + | 3.8 | 4.68E-04 |
| translation (GO:0006412) | 105 | 22 | 5.81 | + | 3.79 | 2.61E-04 |
| nucleoside phosphate biosynthetic process (GO:1901293) | 72 | 15 | 3.98 | + | 3.77 | 2.03E-02 |
| ncRNA processing (GO:0034470) | 73 | 15 | 4.04 | + | 3.71 | 2.34E-02 |
| alpha-amino acid biosynthetic process (GO:1901607) | 85 | 17 | 4.7 | + | 3.62 | 9.63E-03 |
| RNA processing (GO:0006396) | 77 | 15 | 4.26 | + | 3.52 | 4.00E-02 |
| peptide biosynthetic process (GO:0043043) | 113 | 22 | 6.25 | + | 3.52 | 7.74E-04 |
| amide biosynthetic process (GO:0043604) | 140 | 27 | 7.74 | + | 3.49 | 5.23E-05 |
| cellular amino acid metabolic process (GO:0006520) | 190 | 35 | 10.51 | + | 3.33 | 1.49E-06 |
| RNA metabolic process (GO:0016070) | 170 | 31 | 9.4 | + | 3.3 | 1.67E-05 |
| peptide metabolic process (GO:0006518) | 133 | 23 | 7.36 | + | 3.13 | 2.63E-03 |

| | | | | | | |
|---|---|---|---|---|---|---|
| organonitrogen compound biosynthetic process (GO:1901566) | 406 | 68 | 22.46 | + | 3.03 | 3.08E-13 |
| carboxylic acid biosynthetic process (GO:0046394) | 157 | 26 | 8.68 | + | 2.99 | 1.23E-03 |
| organic acid biosynthetic process (GO:0016053) | 159 | 26 | 8.79 | + | 2.96 | 1.52E-03 |
| gene expression (GO:0010467) | 244 | 39 | 13.5 | + | 2.89 | 5.90E-06 |
| cellular amide metabolic process (GO:0043603) | 178 | 28 | 9.85 | + | 2.84 | 1.20E-03 |
| alpha-amino acid metabolic process (GO:1901605) | 138 | 21 | 7.63 | + | 2.75 | 3.83E-02 |
| cellular macromolecule biosynthetic process (GO:0034645) | 178 | 27 | 9.85 | + | 2.74 | 3.44E-03 |
| small molecule biosynthetic process (GO:0044283) | 225 | 34 | 12.45 | + | 2.73 | 1.99E-04 |
| carboxylic acid metabolic process (GO:0019752) | 327 | 48 | 18.09 | + | 2.65 | 1.45E-06 |
| oxoacid metabolic process (GO:0043436) | 334 | 49 | 18.47 | + | 2.65 | 9.46E-07 |
| cellular nitrogen compound biosynthetic process (GO:0044271) | 365 | 53 | 20.19 | + | 2.63 | 2.03E-07 |
| organic acid metabolic process (GO:0006082) | 341 | 49 | 18.86 | + | 2.6 | 1.49E-06 |
| small molecule metabolic process (GO:0044281) | 507 | 70 | 28.04 | + | 2.5 | 7.71E-10 |
| nucleic acid metabolic process (GO:0090304) | 354 | 47 | 19.58 | + | 2.4 | 3.16E-05 |
| cellular component organization or biogenesis (GO:0071840) | 197 | 26 | 10.9 | + | 2.39 | 4.79E-02 |
| cellular biosynthetic process (GO:0044249) | 616 | 81 | 34.07 | + | 2.38 | 7.21E-11 |
| macromolecule biosynthetic process (GO:0009059) | 251 | 33 | 13.88 | + | 2.38 | 6.22E-03 |
| biosynthetic process (GO:0009058) | 650 | 85 | 35.95 | + | 2.36 | 1.44E-11 |
| organic substance biosynthetic process (GO:1901576) | 625 | 81 | 34.57 | + | 2.34 | 1.28E-10 |
| nucleobase-containing compound metabolic process (GO:0006139) | 495 | 64 | 27.38 | + | 2.34 | 1.73E-07 |
| cellular aromatic compound metabolic process (GO:0006725) | 587 | 74 | 32.47 | + | 2.28 | 9.26E-09 |
| heterocycle metabolic process (GO:0046483) | 589 | 74 | 32.58 | + | 2.27 | 1.04E-08 |
| cellular protein metabolic process (GO:0044267) | 255 | 32 | 14.1 | + | 2.27 | 2.65E-02 |
| organic cyclic compound metabolic process (GO:1901360) | 614 | 77 | 33.96 | + | 2.27 | 3.34E-09 |

| | | | | | | |
|---|---|---|---|---|---|---|
| cellular nitrogen compound metabolic process (GO:0034641) | 703 | 86 | 38.88 | + | 2.21 | 3.79E-10 |
| organonitrogen compound metabolic process (GO:1901564) | 781 | 94 | 43.2 | + | 2.18 | 3.34E-11 |
| nitrogen compound metabolic process (GO:0006807) | 1124 | 129 | 62.17 | + | 2.07 | 1.17E-16 |
| cellular macromolecule metabolic process (GO:0044260) | 567 | 64 | 31.36 | + | 2.04 | 3.36E-05 |
| cellular metabolic process (GO:0044237) | 1323 | 144 | 73.18 | + | 1.97 | 7.22E-18 |
| protein metabolic process (GO:0019538) | 383 | 41 | 21.18 | + | 1.94 | 4.45E-02 |
| macromolecule metabolic process (GO:0043170) | 821 | 87 | 45.41 | + | 1.92 | 4.67E-07 |
| primary metabolic process (GO:0044238) | 1277 | 133 | 70.63 | + | 1.88 | 7.50E-14 |
| organic substance metabolic process (GO:0071704) | 1435 | 148 | 79.37 | + | 1.86 | 1.76E-16 |
| metabolic process (GO:0008152) | 1590 | 158 | 87.95 | + | 1.8 | 5.66E-17 |
| cellular process (GO:0009987) | 1918 | 185 | 106.09 | + | 1.74 | 5.42E-23 |
| biological_process (GO:0008150) | 2336 | 198 | 129.21 | + | 1.53 | 3.01E-19 |
| Unclassified (UNCLASSIFIED) | 1768 | 29 | 97.79 | - | 0.3 | 0.00E+00 |

**Supplementary Table 7:** List of eight genes with evidence of positive selection based on the global test

| Gene | (M0)[1] | Likelihood ratio test P-value[2] |
|---|---|---|
| cya | 1.24241 | 0.00669718 |
| group_454 | 2.06375 | 0.00918337 |
| group_1057 | 2.2049 | 0.0029329 |
| group_3049 | 2.49761 | 0.00248336 |
| group_3542 | 1.94168 | 0.00235286 |
| group_3757 | 6.53607 | 0.00434296 |
| group_5674 | 1.58194 | 0.0079647 |
| group_7848 | 9.60069 | 0.00868936 |

[1] M0 estimates a single w across the entire phylogeny of sequences

[2] The p-value of tests after FDR correction. LRT between the global model which estimates omega for each gene and the model which sets omega to 1 for each gene.

**Supplementary Table 8:** GO analysis of accessory genes with significant evidence for positive selection under the sites models M2a vs. M1a

| GO biological process complete | REFLIST | upload | expected | over/under | fold Enrichment | P-value |
|---|---|---|---|---|---|---|
| protein secretion by the type IV secretion system (GO:0030255) | 4 | 4 | 0.01 | + | > 100 | 2.42E-06 |
| secretion by the type IV secretion system (GO:0044097) | 5 | 4 | 0.02 | + | > 100 | 4.35E-06 |
| protein transmembrane transport (GO:0071806) | 40 | 4 | 0.13 | + | 31.57 | 4.41E-03 |
| establishment of protein localization to extracellular region (GO:0035592) | 48 | 4 | 0.15 | + | 26.31 | 8.66E-03 |
| protein secretion (GO:0009306) | 48 | 4 | 0.15 | + | 26.31 | 8.66E-03 |
| protein localization to extracellular region (GO:0071692) | 48 | 4 | 0.15 | + | 26.31 | 8.66E-03 |
| secretion by cell (GO:0032940) | 49 | 4 | 0.16 | + | 25.77 | 9.35E-03 |
| secretion (GO:0046903) | 49 | 4 | 0.16 | + | 25.77 | 9.35E-03 |
| export from cell (GO:0140352) | 51 | 4 | 0.16 | + | 24.76 | 1.09E-02 |
| protein transport (GO:0015031) | 84 | 5 | 0.27 | + | 18.79 | 2.76E-03 |
| protein localization (GO:0008104) | 88 | 5 | 0.28 | + | 17.94 | 3.43E-03 |
| establishment of protein localization (GO:0045184) | 88 | 5 | 0.28 | + | 17.94 | 3.43E-03 |
| macromolecule localization (GO:0033036) | 106 | 5 | 0.34 | + | 14.89 | 8.21E-03 |
| nitrogen compound transport (GO:0071705) | 129 | 6 | 0.41 | + | 14.68 | 9.20E-04 |
| organic substance transport (GO:0071702) | 168 | 6 | 0.53 | + | 11.27 | 4.08E-03 |
| Unclassified (UNCLASSIFIED) | 1768 | 3 | 5.6 | - | 0.54 | 0.00E+00 |

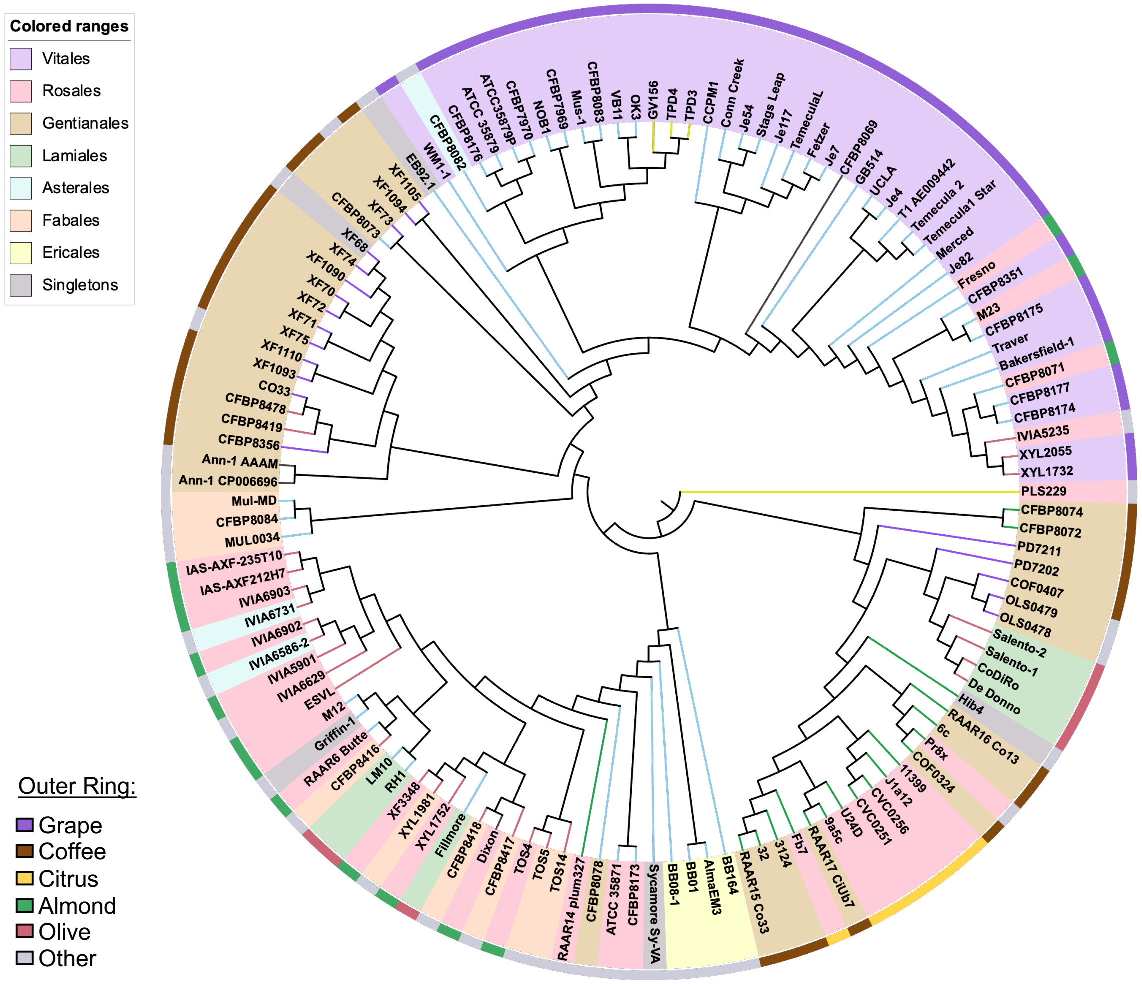


**Supplementary Figure 1:** Phylogenetic relationships of the set of 129 publicly available and novel *Xylella fastidiosa* and *X. taiwanensis* genomes gathered to develop this study. The outer ring denotes the specific plant host from which the bacteria was isolated, and the shaded ranges denote the host plant's taxonomic order. The branches are colored to denote the continent of isolation: North America (Blue), Central America (Purple), South America (Green), Europe (Red), Asia (Gold).

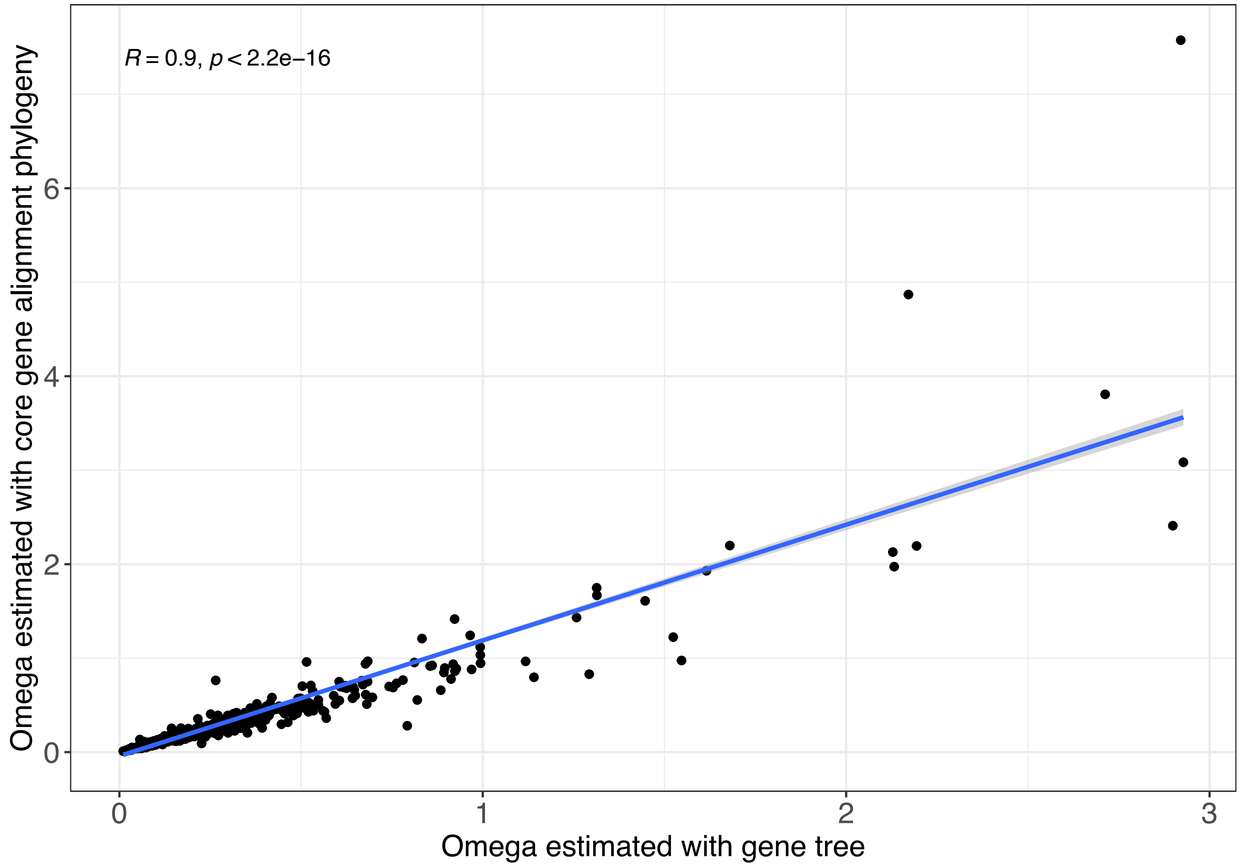


**Supplementary Figure 2:** Correlation plot between the values of omega estimated with two different methods: building an unrooted maximum-likelihood gene tree plotted on the x-axis and with the global phylogeny built from the core gene alignment on the y-axis.

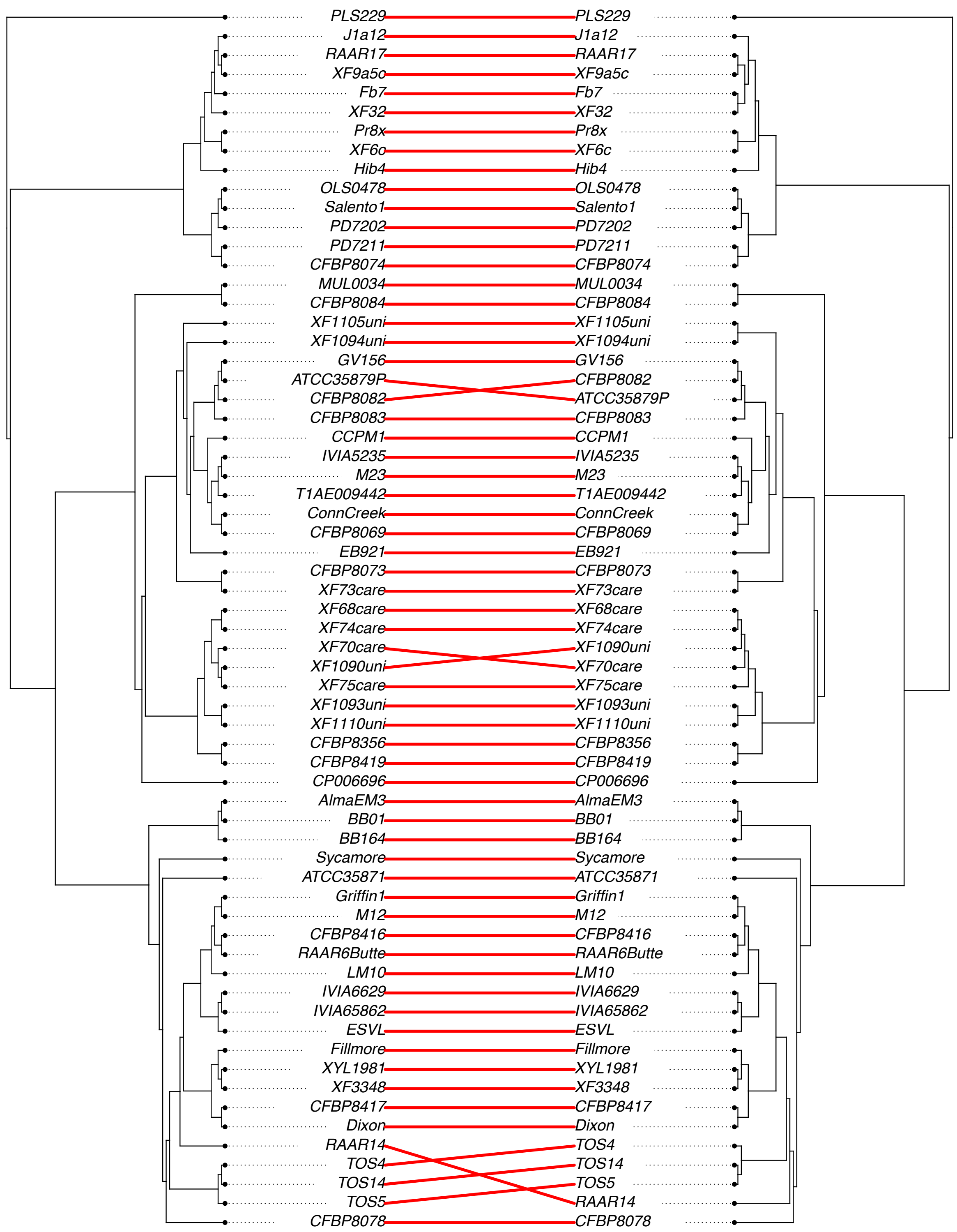


**Supplementary Figure 3:** Comparison of Gubbins recombination-corrected phylogeny (left) to the phylogeny built from the entire core gene alignment (right).

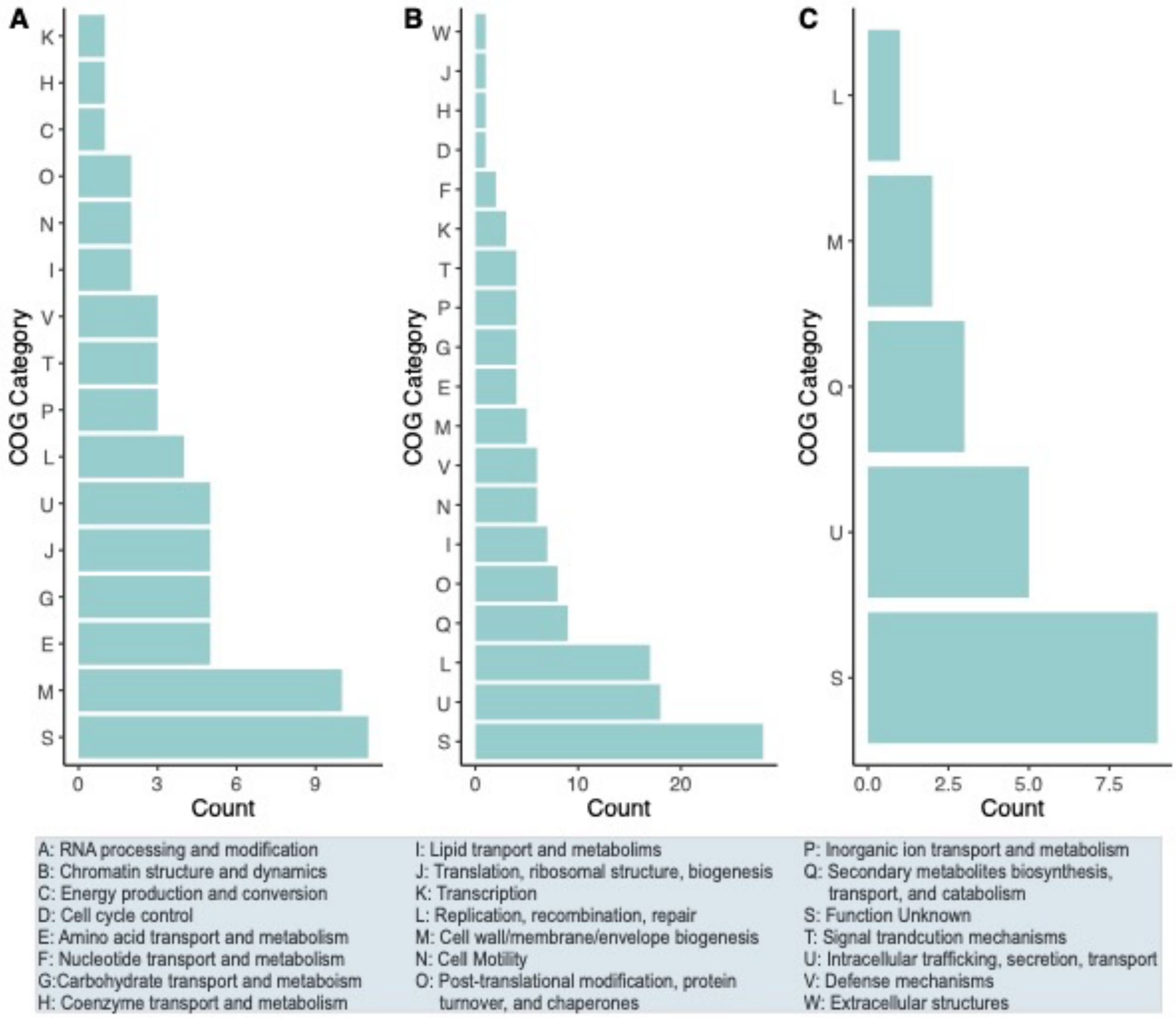


**Supplementary Figure 4:** The distribution of functional categories for A) the 67 core genes, B) the 201 accessory genes, and C) 33 multicopy genes with evidence for positive selection under the sites models. A key to the COG categories is provided in the bottom panel.
